# Distinct flagellins differentially fine tune biofilm initiation via flagellar stator-associated proteins

**DOI:** 10.64898/2025.12.30.697036

**Authors:** Xiaolin Liu, Gunnar N. Eastep, Nataly Torres-Mejia, Ayisha Zia, Karen M. Ottemann, Fengbin Wang

## Abstract

*Helicobacter pylori* relies on flagellar motility to colonize the gastric niche. While *H. pylori* flagella serve multiple functions, including swimming motility, it has recently been appreciated that flagellar basal body properties also contribute to biofilm initiation. The *H. pylori* flagellar filament contains two spatially segregated flagellins, FlaA and FlaB, but their roles in the biofilm initiation process remained undefined. Here, we confirmed that both FlaA and FlaB are required for optimal motility, but found that they exert opposite effects on biofilm initiation: cells without FlaA display low biofilm initiation, while cells without FlaB display elevated initiation. To understand the molecular basis for this divergence, cryo-electron microscopy (cryo-EM) analysis of native flagellar filaments revealed that FlaA and FlaB have distinct supercoiled waveforms. We confirmed these intrinsic waveform curvatures by analyzing filaments from Δ*flaA* and Δ*flaB* mutants, providing a physical basis for their functional specialization. Genetic epistasis experiments further demonstrated that the enhanced early biofilm formation of the Δ*flaB* mutant depends on PilO, a stator-associated component of the flagellar motor recently shown to drive a reciprocal biofilm and motility response. Our findings establish that *H. pylori* has developed functionally specialized flagellins that work with motor-dependent signaling to dynamically balance surface colonization and motility.

## Introduction

Most bacterial species are motile at some point in their life cycle, and they use chemotaxis to move toward or away from nutrients and hostile environments through various forms of movement like swimming, swarming, and twitching (1). The flagellum, a rotary machine, facilitates both swimming and swarming. The extracellular, microns long flagellar filament is built from thousands of copies of flagellins that have a conserved D0/D1 domain architecture across different bacterial species. Beyond enabling movement, flagella are also involved in other critical functions, such as surface adhesion (2, 3), colonization (4), and biofilm formation (5). These additional roles are typically linked to non-conserved features in flagellin, such as the presence of outer domains, the D2, D3, D4 and additional domains (6) or the incorporation of multiple flagellin types that confer additional properties into a single flagellar filament (7).

Biofilm formation is a key survival strategy that allows bacteria to withstand environmental stresses, including antibiotics (8). The relationship between flagellar motility and biofilm formation is often complex and inversely regulated (9). While flagellar motility is essential for the initial stages of surface attachment and biofilm initiation, it is frequently downregulated as the biofilm matures (9, 10). Consequently, mutants lacking functional flagella in diverse bacteria, including *Pseudomonas aeruginosa* (11), *Proteus mirabilis* (12), *Listeria monocytogenes* (5), and *E. coli*/*Salmonella* (13, 14), exhibit significant defects in biofilm formation. This dual role has led to the hypothesis that the flagellum acts as a mechanosensory hub, coupling its motor function to the regulation of surface colonization (15, 16). For instance, in bacteria like *Pseudomonas aeruginosa*, *Vibrio cholerae*, and *Caulobacter crescentus* (17–20), signals from flagellar rotation and assembly directly modulate the production of the extracellular matrix. However, how the flagellin proteins themselves impact biofilm initiation remains poorly understood, particularly in pathogens with unique flagellar adaptations.

*Helicobacter pylori*, a gastric pathogen that infects half the global population and accounts for 75% of the attributable risk for gastric adenocarcinoma cases (21, 22), exemplifies this intricate interplay between flagella and biofilm formation. While flagellar motility is essential for *H. pylori* to colonize the viscous mammalian gastric mucosa (23–25), its flagellar system also regulates biofilm formation in a manner reciprocal to motility. Recent evidence highlights two key mechanisms used by *H. pylori* for this response. First, a cage-like structure at the flagellar base, composed of orthologs of the type four pili proteins PilO, PilN, and PilM, suppresses biofilm initiation and promotes motility (26, 27). Mutants lacking *pilO*, *pilN,* and/or *pilM* display low biofilm initiation and are hyper migratory on semisolid agar (26, 27). Second, the direction of flagellar rotation fine-tunes this balance, with counterclockwise (CCW) rotation promoting biofilm initiation and clockwise (CW) rotation inhibiting it (28). Notably, *H. pylori* lacks all genes encoding proteins related to the common second messenger c-di-GMP, which typically governs the motility-biofilm switch in other bacteria. suggesting a distinct regulatory mechanism (29). These pieces of evidence underscore the flagellum’s dual role as both a propulsion machine and a regulatory scaffold for biofilm development, even in the absence of c-di-GMP.

One unanswered question is how the unusual composition of the *H. pylori* flagellar filament itself impacts biofilm initiation. The *H. pylori* flagellum is sheathed by outermembrane, and composed of two spatially segregated flagellins: FlaB, a minor flagellin located near the hook, and FlaA, the major flagellin located distal to FlaB (30, 31). These two flagellins share 58% sequence identity (32), a much lower similarity than that seen in other microbes, including *Shewanella putrefaciens* (86% sequence identity), which also has spatially segregated flagellins (7). In *S. putrefaciens*, this spatial segregation arises from transcriptional control, and a similar mechanism may take place in *H. pylori*. Specifically, the *H. pylori* gene for the hook-proximal flagellin *flaB* is σ54-dependent. After completion of the hook structure, the anti-σ28 factor FlgM is sequestered, and this allows σ28 to transcribe the *flaA* gene encoding the distal flagellin (7). Transcription of *H. pylori flaA* (σ28-dependent) and *flaB* (σ54-dependent) are known to be differentially regulated, including during biofilm formation (33–35), suggesting the intriguing idea that there may be variation in the amount of these flagellins within flagella. This observation highlights the need for high-resolution structural information and genetic studies to understand their distinct roles.

Here, we investigate the properties of the *H. pylori* FlaA and FlaB flagellins, using a combination of genetic, physiology, imaging, and structural approaches. As reported previously, both FlaA and FlaB are required for optimal motility but to different degrees. Unexpectedly, we discovered they have opposite effects on biofilm initiation: *H. pylori* lacking FlaA are poor biofilm initiators, while those lacking FlaB display elevated early biofilm formation. To uncover the molecular basis for this divergence, we used cryo-electron microscopy (cryo-EM) to reconstruct FlaA and FlaB segments within the wild-type (WT) flagella. These reconstructions revealed that the two flagellins have distinct supercoil waveforms and different surface glycosylation patterns, and exhibit no specific interactions with the sheath detectable by cryo-EM. These intrinsic waveform curvatures were confirmed by analyzing filaments from Δ*flaA* (FlaB only) and Δ*flaB* (FlaA only) mutants, suggesting they represent stable, low-energy conformations for each flagellin. Moreover, by combining our flagellin mutants with a deletion of the flagellar cage protein PilO, we demonstrate that FlaB’s inhibitory effect on biofilm initiation requires PilO. This finding suggests a model in which flagellar filament properties regulate a cellular response, including biofilm initiation, acting through the flagellar basal body and cage. Together, these findings unveil a strategy wherein *H. pylori* employs two functionally specialized flagellins to help the microbe dynamically balance motility and biofilm initiation, a route that likely optimizes survival in the hostile gastric niche.

## Results

### FlaA and FlaB contribute to motility to different extents

To dissect functional roles of the FlaA and FlaB flagellins, we generated Δ*flaA* and Δ*flaB* mutants in the *H. pylori* strain G27. While similar mutations have been characterized in other *H. pylori* strains (23, 31, 32), we first confirmed their key phenotypes in this genetic background, including flagella length, migration in semisolid agar, and swimming in liquid. As expected, the Δ*flaA* mutants exhibited severely truncated flagellar filaments measuring only about 20-40% of WT (Fig. 1A, 1B; Fig. S1). Consistent with this severe truncation, Δ*flaA* mutants were as severely impaired in their migration in semisolid agar as non-motile mutants, with no expanded colony formation (Fig. 1C). The Δ*flaA* strain retained motility in liquid, however, with a speed that was ∼50% that of wild type (WT) (11.4µm/sec vs. 23.6µm/sec) (Fig. 1D). In contrast, Δ*flaB* mutants retained full, WT length flagellar filaments (Fig. 1A, 1B). This mutant was only partially defective in semisolid agar migration (Fig. 1C) and showed just a modest reduction in swimming speed to 20.5µm/sec (Fig. 1D). We also measured the direction-change (reversal) frequency, which reflects the transition from counterclockwise (CCW) to clockwise (CW) rotation (36). While Δ*flaB* mutants displayed reversal frequencies comparable to WT, Δ*flaA* mutants showed a significantly lower frequency of direction changes (Fig. S1). Together, these results demonstrate that both FlaA and FlaB are necessary for optimal *H. pylori* motility, but to different degrees. Deletion of *flaA* results in significantly shortened flagella, impaired motility, and CCW-prone flagellar rotation bias. In contrast, deletion of *flaB* results in strains that have normal length flagella and only minimal effects on motility.

**Figure 1.**
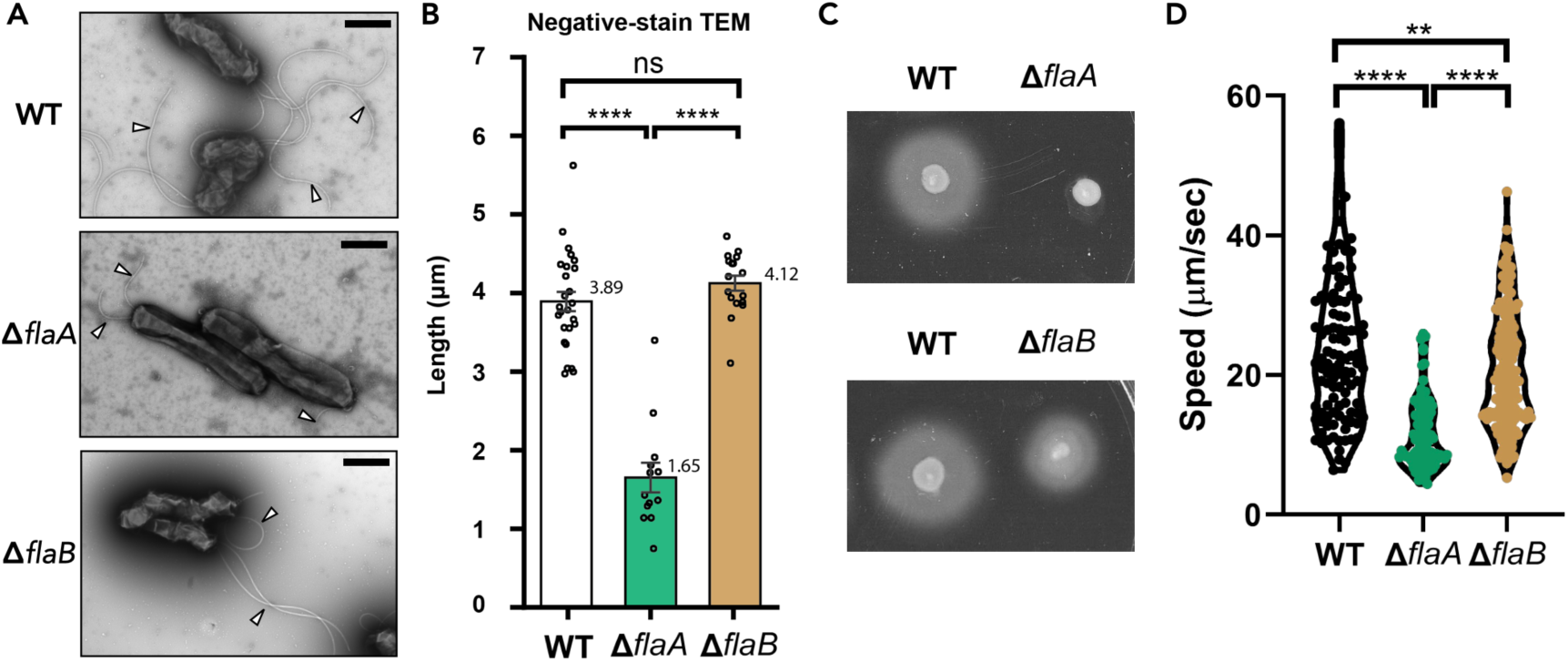
FlaA plays a major role in flagellation and motility while FlaB plays only a minor role. *H. pylori* strain G27 was deleted for *flaA* or *flaB* and analyzed for flagellation and motility. (A) Negative staining images show that WT, Δ*flaA*, and Δ*flaB* cells produce flagella when grown in liquid. Observed supercoiled flagellar filaments are indicated with white arrowheads. The scale bar indicates 1 μm. (B) Flagella lengths measured from negative stain transmission electron microscopy images of WT, Δ*flaA*, and Δ*flaB* cells. Bars represent the mean ± standard error of the mean. Statistical significance was determined by ANOVA and Tukey’s HSD multiple comparisons of the mean; **** p < 0.0001. (C) Spreading diameter of *H. pylori* WT, Δ*flaA*, and Δ*flaB* in 0.35% (w/v) semisolid agar after culturing for 3 days. (D) Swimming speeds of *H. pylori* WT, Δ*flaA*, and Δ*flaB* cells. Overnight liquid cell cultures were adjusted to OD_600_ of 0.15 and recultured in fresh BB10 media for 2 hours. The swimming trajectory of individual cells was recorded under phase-contrast microscope. The speed of each cell was calculated using ImageJ.

### FlaA and FlaB have antagonistic impacts on biofilm initiation

Several studies have shown that *H. pylori* can form biofilms on both biotic and non-biotic surfaces (37–42), a property known to be affected in other microbes by flagellar components (26, 28, 42). To investigate the specific roles of *H. pylori* FlaA and FlaB in this process, we characterized the biofilm phenotypes of Δ*flaA* and Δ*flaB* mutants on two surfaces, polystyrene and glass, using two distinct methods of monitoring biofilm creation. *H. pylori* G27 WT begins to form biofilm after one day of incubation, the initiation phase, and reaches maturation after three days (28, 34). During the biofilm initiation phase (day 1), the Δ*flaA* and Δ*flaB* strains exhibited statistically significant and opposing differences in biofilm mass compared to WT (Fig. 2A). The Δ*flaA* strain produced approximately 5-fold less biofilm mass than WT; we note that this reduction likely reflects both the significant truncation of the flagellar length in this mutant and decreased motility. In contrast, the Δ*flaB* strain produced 2-fold more biofilm mass at day 1 (Fig. 2A). This hyper-biofilm phenotype came as a surprise, as it occurs in the context of a full-length flagella and near-normal motility (Fig. 1). This result thus suggested the idea that FlaB acts normally to suppress biofilm initiation independently of gross defects in flagellar structure or motility. By day 3, the total biomass of mature biofilms was not significantly affected by the loss of either *flaA* or *flaB* (Fig. 2B). These results suggest that FlaA and FlaB play critical and antagonistic roles during the early biofilm initiation stages.

**Figure 2.**
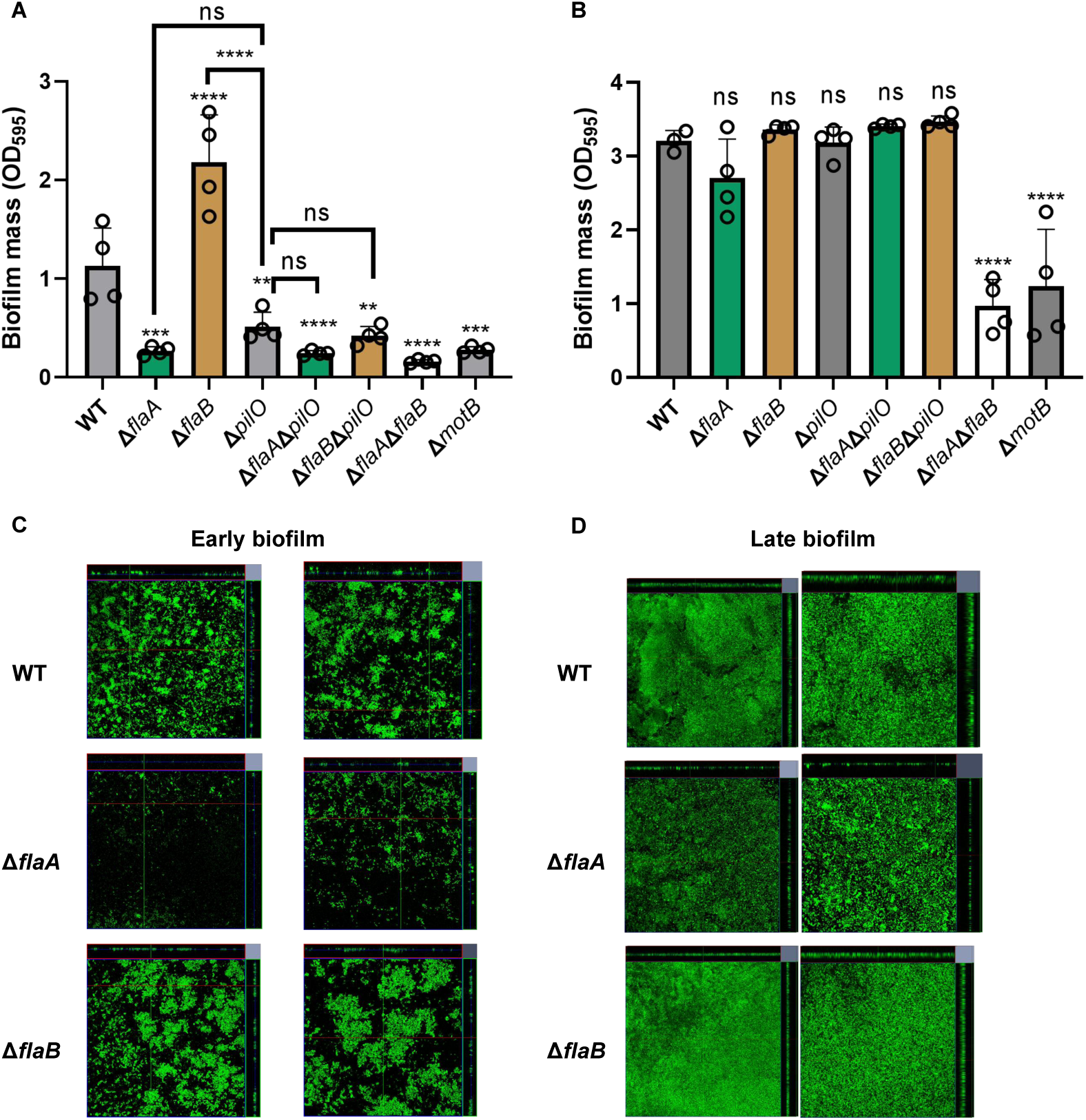
FlaA and FlaB act antagonistically to either activate or suppress biofilm initiation. Biofilms were allowed to form under static conditions in microtiter plates for 1 day (A) or 3 days (B). Overnight liquid cell cultures were set to OD_600_ of 0.15 with fresh BB10 media and then transferred into 96-well plate to develop biofilm. Biofilm formed on the well surface was stained with 0.1% (w/v) crystal violet for 10 min and quantified by OD_595_. The data are presented as the mean of at least three biological replicates, with each replicate shown as an open circle. Error bars are SEs. The asterisks indicate a significant pairwise difference between strains according to Student’s t test (**P < 0.0015; ***P < 0.001;****P < 0.0001; ns, no significant difference). Representative images of GFP-expressing WT, Δ*flaA*, and Δ*flaB* biofilms after culturing for 1 or 3 days under static conditions on glass coverslips (C-D) were imaged using confocal microcopy. (C) After 1 day, WT cells attached singly or in small clusters. Δ*flaA* mutants showed reduced attachment primarily as single cells, while Δ*flaB* mutants exhibited increased clustering and larger microcolonies. (D) After 3 days, all strains formed thick, multi-layered biofilms. WT biofilms displayed a patchy structure with variable thickness; Δ*flaB* biofilms were more uniform, while Δ*flaA* biofilms showed substantial surface unevenness. The large central panels show top-down views; adjacent side panels display vertical cross-sections taken at the indicated lines.

To support these findings, biofilms were formed on glass slides and imaged using confocal laser scanning microscopy of GFP-expressing WT, Δ*flaA,* or Δ*flaB* strains. After one day, the WT strain formed attached cells that were either single or in small clusters (Fig. 2C). Compared to WT, the Δ*flaA* mutant had fewer attached cells, which were predominantly found as single, isolated cells. In contrast, the Δ*flaB* strain exhibited high numbers of large, densely packed cell clusters, that appeared more substantial than those formed by WT (Fig. 2C). These results, thus, support the crystal violet biofilm findings. After three days, all three strains formed mature, thick, multi-layers consistent with previous descriptions, as predicted from the crystal violet staining (Fig. 2D) (34, 43). Collectively, these results demonstrate that FlaA and FlaB have impactful roles during the initiation of *H. pylori* biofilm but a minor influence on the final mature biofilm architecture. Moreover, these two flagellins act antagonistically: the loss of FlaA decreases early biofilm formation, while the loss of FlaB enhances it.

### Cryo-EM reveals structural differences between FlaA and FlaB in the WT flagellum

To understand the structural basis for the opposing effects of FlaA and FlaB on biofilm formation, we investigated their high-resolution structures using cryo-EM. For this analysis, we isolated flagellar filaments by mechanically shearing them from *H. pylori* WT cells and subsequently removing the cells via centrifugation. Under cryo-EM, this treatment led to the presence of both sheathed (10%) and unsheathed (90%) flagella (Fig. S2). Unsheathed flagella were presumably introduced by the shear force. To reach a high resolution, helical reconstruction was first performed on all unsheathed flagella particles after indexing the average power spectrum from aligned particles. A 360-pixel small box and canonical bacterial flagellum symmetry were used. With over two million particles, the refinement converged at 2.8 Å (Figure 3A-B), with optimized helical rise and twist parameters of 4.77 Å and 65.40 degrees, respectively. The flagellin backbone was clearly traceable, with the AlphaFold (44) model of the major flagellin, FlaA, fitting into the map and no indication of disordered surface residues. Using the unsheathed FlaA structure as a reference, the map was low-pass filtered to 20 Å and used as the reference to reconstruct the sheathed flagella particles. The resulting sheathed flagellum structure reached 4.2 Å resolution, and the flagellin part was identical to the unsheathed one, with no extensive contacts observed between the flagellum and lipids after applying helical symmetry (Fig. S3). Both 2D averages and 3D reconstructions of sheathed flagella revealed a lipid bilayer structure. The peak-to-peak distance within the bilayer, measured from the cylindrically averaged 3D map, was approximately 36 Å, consistent with the LPS structure (Fig. S3) (45).

**Figure 3.**
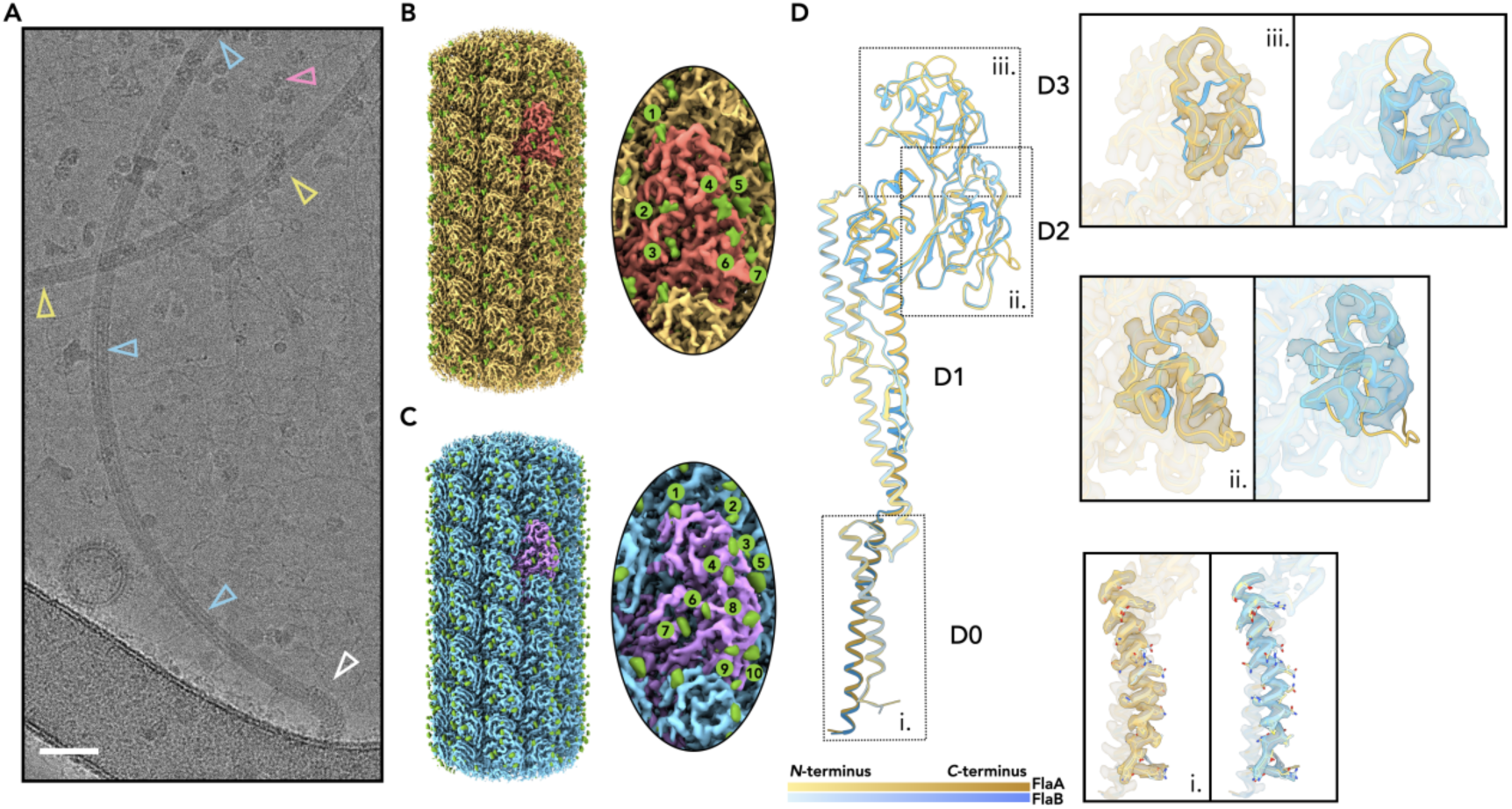
Cryo-EM of *H. pylori* flagellar filaments reveal the spatial distribution of flagellins. (A) A representative cryo-EM micrograph of sheared wild-type *H. pylori* flagellar filaments. The observed flagellar hook is indicated by a white arrowhead. FlaB particles are indicated by blue arrowheads. The remaining flagellar filaments, lacking a visible hook, were assumed to be predominantly FlaA regions and used for FlaA reconstruction (yellow arrowheads). Observed ureases are indicated by a pink arrowhead. Scale bar: 50 nm. (B) Helical reconstruction of the FlaA region of *H. pylori* flagella. FlaA are colored yellow, with one copy highlighted in red. Extra densities corresponding to monosaccharides, are colored green. A zoomed-in view of glycosylation is shown on the right. (C) Similar to (B), a helical reconstruction of the FlaB region is shown, with FlaB flagellin colored cyan and one copy highlighted in purple. Extra cryo-EM densities, corresponding to monosaccharides, are colored green. A zoomed-in view of glycosylation is shown on the right. (D) The atomic models of FlaA and FlaB, along with the cryo-EM maps, clearly distinguish them. FlaA and FlaB, and their corresponding cryo-EM maps, are colored yellow and cyan, respectively, as in panels B and C. For the atomic models, the color is graded from the N-terminus to the C-terminus. Three different regions are zoomed in on the right: FlaA and FlaB maps are displayed separately, with both models docked to clearly demonstrate their differences.

To define the structure of FlaB within the WT flagella, we took advantage of the fact that it forms a hook-proximal segment of ∼1-2 μm in length in *H. pylori* (31). Accordingly, we manually selected micrographs with visible hook structures. Filaments were automatically picked from these micrographs using the CryoSPARC ‘Filament Tracer’, removing any that lacked hooks. Hook-connected segments were chosen as within the 1 μm of the hook end. In this way, ∼220K FlaB particles were selected, and used to create a 3.2 Å resolution in the helical reconstruction (Fig. 3C), judged by the map:map FSC (Fig. S4). The sequence of FlaB, but not FlaA, can be threaded into the cryo-EM map with great confidence. The sequence identities of the two flagellins is 59.8%, with the greatest disparity in the D2/D3 domains (Fig. S5). Visual inspection of the two maps and models (Fig. 3D) reveal segments where the maps and models of FlaA clearly diverge from the FlaB map and model. Aside from tertiary-structural differences, the two proteins also differ in their glycosylation. Early mass-spectrometry studies of *H. pylori* flagellins ascertained that FlaA is glycosylated at seven sites and FlaB at ten sites by the sugar pseudaminic acid (46), and all of them were clearly observed in the cryo-EM maps (Fig. 3B-C, Fig. S6). Overall, these experiments support that the FlaB is hook-proximal and the two flagellins can be structurally delineated using position.

*flaB* is encoded in the *H. pylori* genome in an operon with the *pseE* glycosyltransferase downstream (47, 48) We thus wanted to validate that the antibiotic resistance gene cassette in the Δ*flaB* mutant does not have polar effects on *pseE*, which might indirectly affect biofilm formation by altering FlaA glycosylation, although this possibility was unlikely as previous work showed that FlaA cannot assemble into the flagella without glycosylation (47, 48). We therefore performed a direct comparative analysis of glycosylation densities using our high-resolution cryo-EM data compared FlaA in WT cells to that in the Δ*flaB* mutant (FlaA only). The helical reconstructions of the wild-type FlaA filament and the FlaA-only filament reached the same resolution of 2.79 Å (Fig. S7). The glycosylation sites exhibited no visible differences between the two. To compare the glycosylation densities, both maps were sharpened using the same B-factor (-80 Å^2^). Next, the map threshold were normalized to each other based on the density of the ordered protein domains (D0/D1 region). Subsequently, when we compared the glycosylation densities of both maps at different display thresholds (Fig. S7), we observed that both the glycan and protein densities were virtually identical between the wild-type and FlaA-only filaments, indicating no changes in the glycosylation level of FlaA between the two. This analysis confirms that PseE function remains intact in the Δ*flaB* mutant, and that the hyper-biofilm phenotype is a direct consequence of losing the FlaB flagellin.

### FlaA and FlaB exhibit distinct supercoiled waveforms

Beyond D2/D3 domain-level structural differences, we observed divergent supercoiling properties between FlaA- and FlaB-dominated filament segments within native flagella. Using cryoSPARC’s Filament Tracer, we quantified 2D curvature in WT filaments: FlaA-rich distal regions showed lower curvature (κ ≈ 1.7 μm^-^¹) compared to FlaB-rich hook-proximal regions (κ ≈ 2.5 μm^-^¹), suggesting distinct waveforms along the native filament (Fig. 4A). To test if these curvatures represent intrinsic, low-energy states for each flagellin, we performed additional cryo-EM analysis on homogeneous filaments from Δ*flaB* (FlaA only) and Δ*flaA* (FlaB only) mutants. This analysis confirmed that the isolated filaments maintained their distinct curvatures: FlaA filaments had a lower curvature (∼1.7 μm^-^¹), while FlaB filaments adopted higher curvature (∼2.5 μm^-^¹), consistently matching their respective regions in WT flagellum (Fig. 4A). We further resolved supercoiled waveform differences through asymmetric reconstructions using extended particle boxes (640-pixel) to accommodate curvature. Final C1 refinements without imposed symmetry yielded 3.0 Å (FlaA) and 3.4 Å (FlaB) resolution maps (Fig. 4B). Full-length supercoiled models generated from atomic structures revealed key geometric differences: FlaA filaments form longer-pitch, smaller-diameter left-handed supercoils compared to FlaB (Fig. 4C).

**Figure 4.**
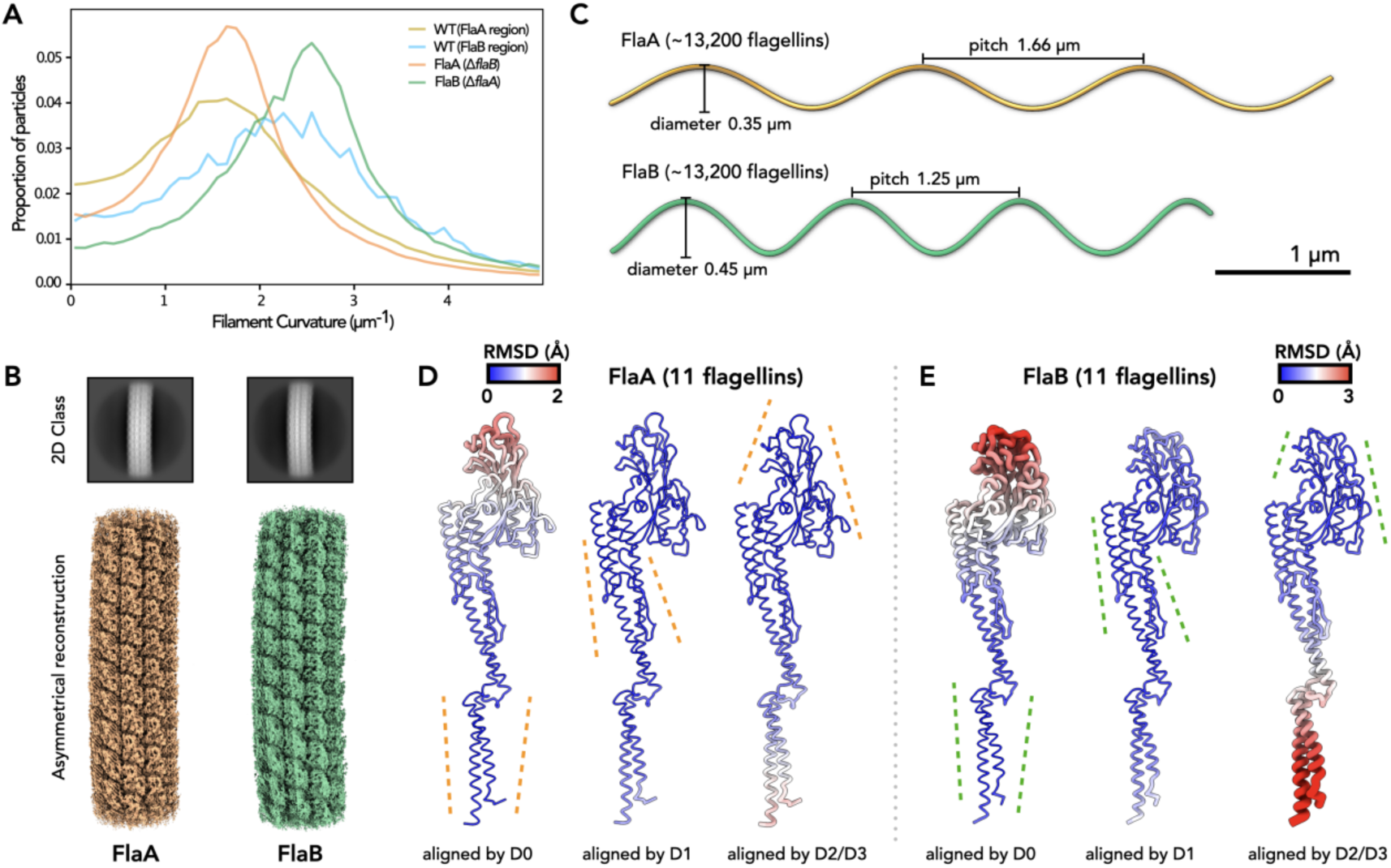
The waveforms of FlaA and FlaB filaments are different. (A) Filament local curvature was estimated using the ‘Filament Tracer’ job in cryoSPARC. Four datasets, including particles used in the final reconstruction, were analyzed: the FlaA region in wild-type flagella, the FlaB region in wild-type flagella, FlaA-only flagella, and FlaB-only flagella. (B) Class 2D averages (640-pixel box) and the final asymmetric reconstructions of FlaA (orange), using the FlaA-only flagella dataset, and FlaB (green), using the FlaB-only flagella dataset. (C) FlaA and FlaB supercoiled model, generated from cryo-EM atomic models from asymmetric reconstructions. This process began with an atomic model containing 55 flagellins (five layers of 11 flagellins, representing 11 protofilaments). The flagellins within each protofilament were treated as identical subunits. The supercoiled filaments were then expanded by adding another 299 atomic models, each containing 55 flagellins. The top layer of one model was aligned to the bottom layer of its adjacent model, resulting in a supercoiled filament with ∼1,200 layers and 13,200 flagellins. To conserve memory, instead of aligning entire layers of 11 flagellins, three atoms per flagellin (one each in the D0, D1, and D2/D3 domains) were selected for alignment and display. (D-E) Comparison of flagellins in all eleven protofilaments. FlaA (D) and FlaB (E) flagellins were aligned by their D0, D1, and D2/D3 domains separately. The RMSD between the 11 flagellins after each alignment is shown using a color gradient. Larger RMSD values correspond to larger cartoon radii.

To understand how FlaA and FlaB assemble into distinct supercoiled waveforms despite high similarity, we examined their conformational flexibility. Previous studies of both straight flagella containing mutated flagellins (49) and wild-type supercoiled flagella (50) indicate flagellin conformation is mediated by the hinge region, a linker connecting the D0 and D1 domains, while the D0 and D1 domains themselves remain almost rigid bodies across different conformations. Consistent with this, alignment of individual domains (D0, D1, or D2/D3) from all eleven FlaA or FlaB conformations revealed minimal RMSD within each domain (Fig. 4D-E), confirming their rigidity. Conformational differences arose instead from rotations within the hinge regions, altering domain-to-domain orientations across protofilaments (Fig. 4D-E). Crucially, the D0-D1 hinge sequence is identical in FlaA and FlaB, eliminating hinge sequence variation as the cause of their distinct waveforms. Instead, we propose that residues involved in inter-flagellin interfaces dictate supercoiling preference, while the hinge enables necessary rotational freedom. Supporting this, a structure-based sequence alignment reveals that many of the non-conserved residues between FlaA and FlaB are located at two major inter-subunit interfaces: the left-handed 5-start and 11-start interface (Fig. S5). In addition, PISA analysis (51) showed FlaB interfaces have a 3% larger buried surface area (6,200 Å² vs. 6,023 Å²) and 13% stronger binding energy (-51.3 kcal/mol vs. -45.2 kcal/mol) than FlaA (Fig. S8). These biophysical differences likely constrain hinge rotation during assembly, ultimately determining the divergent supercoiled waveforms of FlaA and FlaB filaments.

### Δ*flaB* mutants form clustered biofilms with connecting filaments, while Δ*flaA* biofilms are sparse biofilms and lack visible connections

To further investigate the role of flagellins in *H. pylori* biofilm architecture, we switched to scanning electron microscopy (SEM). This technique provides a magnification intermediate between light microscopy and cryo-EM, allowing observation of single cells, flagella, and cell-cell connections. Biofilms were formed on glass coverslips for one or three days. After one day, the WT strain attached to the surface as both rod and coccoid shaped cells, with numerous flagellar filaments connecting cells to each other and to the surface (Fig. 5). Flagella often lose their native supercoiled shape during SEM sample preparation (52), presumably due to drying and metal shadowing; therefore, we separately confirmed the presence of these supercoiled filaments connecting cells using confocal microscopy and the membrane stain FM4-64 (Fig. S9). After three days, the *H. pylori* WT strain formed thick biofilms with a what appears to be a copious extracellular matrix (Fig. S10).

**Figure 5.**
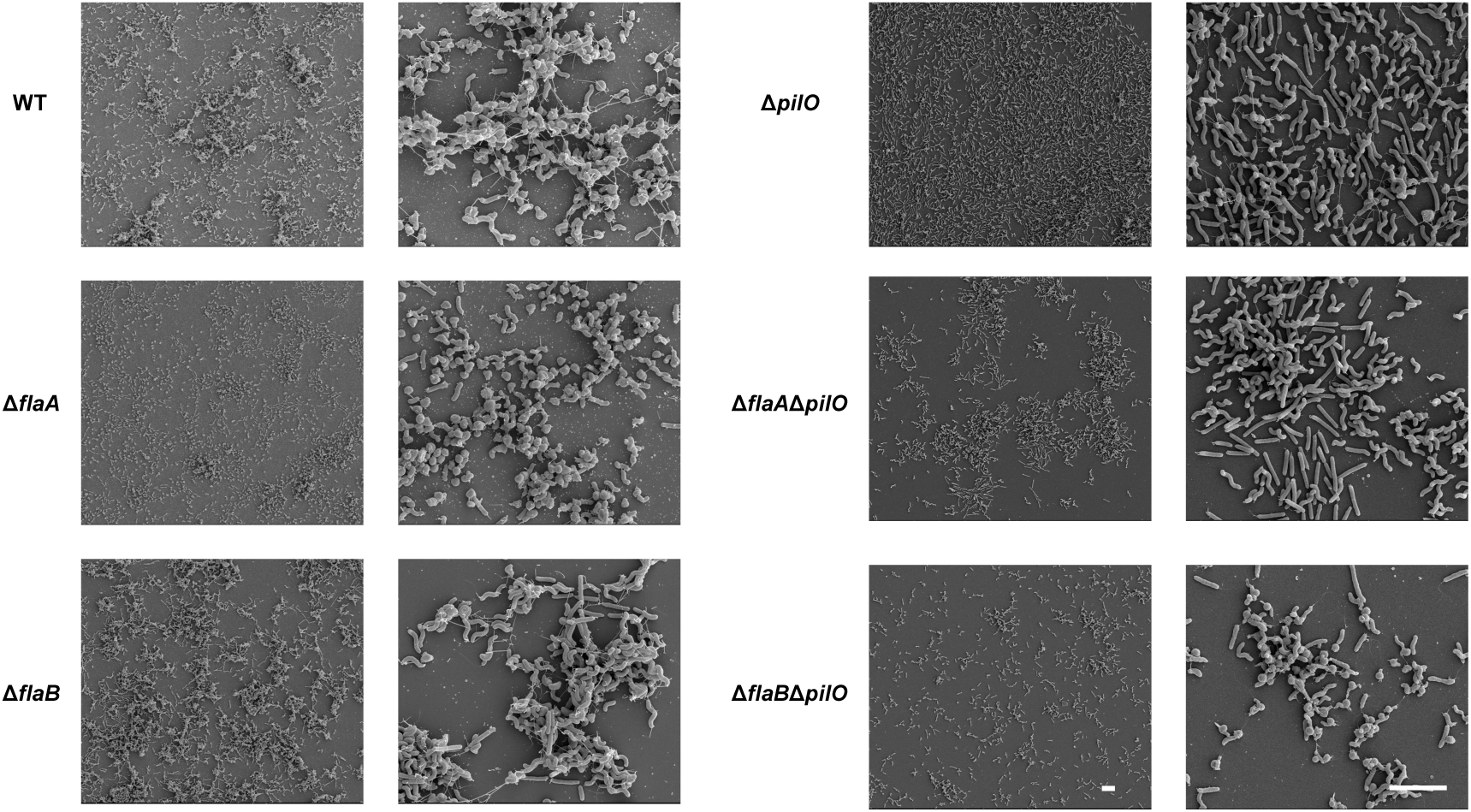
Distinct biofilm architectures of *flaA* and *flaB* mutants reflect flagellar phenotypes, and PilO is required for biofilm cluster formation and mediates FlaB-dependent biofilm suppression. Scanning electron micrographs (SEM) of *H. pylori* G27 wild-type (WT), Δ*flaA*, Δ*flaB*, Δ*pilO*, Δ*flaA*Δ*pilO* and Δ*flaB*Δ*pilO* cells have flagella after culturing on glass surface for 1 day. WT biofilms show attached rod and coccoid cells interconnected by numerous flagellar filaments. Δ*flaB* mutants form larger cell clusters than WT, with flagellar filaments attaching to surfaces and connecting cells. Δ*flaA* mutants exhibit sparse attachment with truncated flagella unable to bridge neighboring cells. Δ*pilO* mutants possess wild-type-like flagella capable of surface attachment and intercellular bridging, but form uniformly dispersed single cells without clusters. Δ*flaB*Δ*pilO* double mutants phenocopy Δ*pilO* single mutants, exhibiting dispersed single-cell attachment and lacking cell clusters. Δ*flaA*Δ*pilO* mutant have low biofilm capacity. Scale bars: 5 µm. Biofilm samples were fixed, dehydrated, critical point dried, sputter-coated with gold, and imaged at 5 kV as described in Methods.

At day one, Δ*flaB* mutants formed cell clusters that were visibly larger than those of the WT (Fig. 5), which aligns with our results from crystal violet staining and confocal microscopy (Fig. 2). Similar to WT, the Δ*flaB* biofilms also displayed numerous flagellar filaments attaching to the surface and connecting cells (Fig. 5A). At day three, there were no visible differences between WT and Δ*flaB* biofilms; both displayed extensive cell clustering and abundant extracellular matrix-like substance (Fig. S10). In contrast, Δ*flaA* mutants, with their short flagella, exhibited only sparse surface attachment at day one, and their truncated filaments were rarely observed bridging neighboring cells (Fig. 5). By day three, despite accumulating a total biomass comparable to the WT based on crystal violet measurements (Fig. 2), the Δ*flaA* mutant displayed a distinct architecture of rough biofilms with many cells arranged in rope-like structures via direct cell-cell contact (Fig. S10). These results confirm that *H. pylori* lacking *flaB* form more abundant biofilms than WT, and these display high levels of clustering and clumping of cells.

### The biofilm-regulating role of FlaB requires the PilO flagellar cage

Our results show that Δ*flaB* mutants produce elevated levels of biofilm despite having nearly normal flagella. Previously, we established that an inner-membrane protein cage in the flagellar basal body, formed by PilO, PilN and PilM, couples flagellar properties to biofilm regulation (28). We therefore determined if FlaB’s inhibitory role in biofilm initiation depended on the PilM/N/O cage. Δ*pilO* mutants possess functional flagella and exhibit motility equal or better than WT, but they are defective for biofilm initiation (28). SEM analysis confirmed this phenotype, with Δ*pilO* mutants displaying defects in the initial clustering of cells at the biofilm initiation stage (Fig. 5). This observation suggests that functional flagella alone are insufficient for biofilm initiation and cell clustering.

To directly test the genetic interaction between FlaB and PilO, we generated a Δ*pilO*Δ*flaB* double mutant. Strikingly, the Δ*pilO*Δ*flaB* mutant phenocopied the Δ*pilO* single mutant, failing to form the large cell clusters characteristic of the Δ*flaB* strain during early biofilm formation (Fig. 5). Crystal violet assays confirmed this observation, showing that Δ*pilO*Δ*flaB* mutant produced low early biofilm mass similar to that of the Δ*pilO* mutant (Fig. 2). The loss of biofilm initiation indicates that the ‘hyper-biofilm’ potential of the Δ*flaB* mutation cannot be realized without functional PilO. This epistatic relationship, where the double mutant phenocopies the Δ*pilO* single mutant, positions PilO downstream of FlaB in the biofilm pathway. As controls, a Δ*motB* mutant with paralyzed flagella and a Δ*flaA*Δ*flaB* double mutant with no flagella both exhibited severely reduced biofilm after one or three days (Fig. S11). These controls confirm that both the presence and rotation of the flagellum are fundamental requirements for initiating a biofilm. Together, these results demonstrate that the biofilm-suppressing function of the FlaB flagellin is strictly dependent on signaling through the PilO flagellar cage.

## Discussion

In this study, we report the finding that *H. pylori* flagellins have shared functions for promoting motility, but opposite functions with regard to promoting biofilm initiation. In line with this biofilm-based difference, these two flagellins have distinct structures. This latter information arose from a near-atomic resolution cryo-EM structures of the two flagellar filaments, FlaA and FlaB, within the wild-type *H. pylori* flagellum, demonstrating the two proteins exhibit distinct surface glycosylation patterns and form different supercoiled waveforms. For swimming, FlaA plays a dominant role and FlaB a more minor one, but both flagellins are required for optimal performance. A key finding arose from the Δ*flaB* mutant. This mutant possesses WT-length flagella composed of only FlaA and near-normal motility, but forms significantly more biofilm than WT. This result uncouples the biofilm phenotype from general defects in flagellar length or motility. It supports a model in which the presence of FlaB in the WT filament suppresses a pro-biofilm activity associated with FlaA. In contrast, the Δ*flaA* mutant, with its short, FlaB-only filaments, is impaired in both motility and biofilm, a confounded phenotype that underscores the dominant role of FlaA in filament assembly and propulsion. This functional divergence arises despite relatively high sequence similarity between FlaA and FlaB, but of specific differences—distinct spatial positioning, different inter-subunit interfaces, and unique filament curvatures—may underlie this phenotype. Furthermore, we establish that FlaB’s biofilm-suppressing role requires the motor-associated protein PilO, a key component of the flagellar cage that was previously shown to transduce flagellar-based signals into reciprocal biofilm and motility regulation (26, 28).

From a bioenergetic perspective, flagella are a significant investment for bacteria, costing an estimated 0.5–40% of their total energy budget (53). Filaments composed of multiple flagellins present additional evolutionary trade-offs, including higher costs for synthesis and regulation, as well as greater immunological risks. Despite this, complex flagellar systems are surprisingly common. A previous analysis of 607 bacterial genomes in KEGG database revealed that 45% possess multiple flagellin genes (54). This multiplicity arises in two main ways: some bacteria, such as *Azospirillum brasilense* (55), encode multiple distinct flagellar systems (e.g., polar and lateral), while many others, including *H. pylori*, *Shewanella* (7, 56, 57), *Campylobacter jejuni* (58), and *Vibrionaceae* (59), incorporate two or more flagellin types into a single filament. A common feature of these latter genera is their adaptation for swimming in viscous environments, where our work and previous studies have shown that a composite filament is required for optimal motility (7).

What is the molecular basis for this enhanced motility? A compelling hypothesis is that using multiple flagellins creates a gradient of curvature along the filament, improving torque transmission and propulsive efficiency compared to a uniform filament. We can use a car’s drivetrain as an analogy, where a universal joint (the hook) connects the engine (flagellar motor) to a driveshaft (the minor, hook-proximal filament) and then to the wheels (the major, distal filament). This system provides both the rigidity needed for propulsion and the flexibility to handle rotational stress. This argument is strongly supported by the calculated curvatures of these components based on observed pitch and diameter (60). The flagellar hook has a very high curvature (22–28.5 μm⁻¹), in reported structures from *S. enterica* (61) and *S. oneidensis* (56). The major, distal filament, in contrast, has a much lower curvature (∼1.7 μm⁻¹), a value that is remarkably similar across *S. enterica* (61), *S. oneidensis* (56), and FlaA in *H. pylori*. Critically, the minor flagellar filament that connects them has an intermediate curvature of ∼2.5–2.6 μm⁻¹, as measured by cryo-EM in this study and by fluorescence microscopy in *S. oneidensis* (56), perfectly bridging the gap between the hook and the main filament.

The advantages of possessing multiple flagellins extend beyond simply improving motility in complex environments. Emerging evidence suggests that different flagellin proteins can confer different functions (7, 62–64). In *Vibrio anguillarum*, for example, individual flagellin mutants show only minor defects in motility, but have substantial defects in animal colonization (64). In another case, seminal studies on *S. putrefaciens* suggested its two flagellins have specialized mechanical roles: the hook-proximal flagellin provides stability, while the hook-distal flagellin enables the filament to wrap around the cell body for screw-like propulsion (7, 65). It is unclear whether a filament-wrapping mechanism plays a role in *H. pylori* motility. Instead, our work dissecting *flaA* and *flaB* mutants reveals a clear functional specialization related to lifestyle switching in *H. pylori*. We showed that the two flagellins have contrasting roles in biofilm formation: FlaA promotes biofilm initiation and cell-cell bridging, whereas the short filament formed by FlaB alone are ineffective for these functions (Figs. 2, 5). This phenotype was unmasked by using FlaA-only filaments, which formed high levels of biofilm at the early phase. This specialization aligns with their distinct genetic regulation; *flaB* is σ54-dependent, while *flaA* is σ28-dependent (33, 66). Furthermore, σ54 and σ28 expression diverges during biofilm formation (67). Our previous transcriptomics analysis (34) and qRT-PCR analysis (Fig. S12) showed that *flaA* expression increased in biofilm compared to planktonic cells. This suggests *H. pylori* may actively fine-tune biofilm development by modulating the ratio of FlaA and FlaB in its flagellar filaments.

How does the FlaB flagellin inhibit biofilm initiation? Our data support a model where FlaB’s function requires a downstream signaling pathway component, PilO. We hypothesize that the distinct physical properties of the FlaB filament segment, potentially its higher curvature, influence the basal body/cage complex in a manner that is sensed by PilO, leading to the suppression of biofilm initiation programs. The requirement for FlaA is clear, as the high-biofilm phenotype of the Δ*flaB* mutant only occurs when the FlaA filament is present, suggesting FlaA likely provides a pro-biofilm signal that FlaB is positioned to counteract. The second critical component is PilO, which forms an inner membrane cage at the flagella base(68, 69). While deleting *flaB* alone boosts biofilm, combining it with a *pilO* deletion (Δ*flaB*Δ*pilO*) reverts the phenotype to the low-biofilm state of a Δ*pilO* single mutant. This epistatic relationship supports the idea that PilO acts downstream of FlaB in the same regulatory pathway. To date, bacterial flagella have been well-known to function as adhesins and mechanosensors (15, 70–72). While both Δ*flaA* and Δ*flaB* mutants exhibit impaired motility, only the Δ*flaB* mutant displays enhanced biofilm initiation (Fig. 2-5). This finding suggests FlaB’s inhibitory role stems from functions distinct from motility, potentially involving adhesion or mechanosignaling. An alternative, non-exclusive model is that the spatial segregation of FlaA and FlaB could differentially impact the localization of adhesins to the flagellar sheath. For instance, if FlaA recruits specific adhesins (e.g. HpaA (73)), the section of filament formed by FlaB in the wild-type may act as a buffer zone that limits the rate at which adhesins are recruited to the sheath, which might account for some of the enhanced biofilm initiation observed in the Δ*flaB* mutant. Therefore, although our genetic data showing that FlaB’s effect requires the inner membrane protein PilO strongly suggests a signaling model, we cannot fully exclude the contribution of other alternative possibilities, such as this direct adhesion model. Our previous work indicated that *H. pylori* flagellar rotation itself acts as a signal influencing biofilm initiation, with counter-clockwise rotation leading to elevated biofilm formation (28). Δ*flaB* mutants, however, did not alter rotation compared to WT. Our data furthermore argue against a primary role for FlaB-mediated adhesion: the flagella of Δ*flaB*Δ*pilO* double mutant retained normal assembly, motility, and apparent surface attaching abilities based on SEM images, yet exhibited a severe biofilm initiation defect phenocopying the Δ*pilO* mutant. Given PilO’s position as an inner membrane protein associated with the flagellar motor cage, these findings support a model wherein FlaB limits biofilm initiation by facilitating a PilO-dependent signaling pathway, perhaps responsive to external mechanical stimuli, rather than through direct adhesion.

In summary, the cryo-EM structures of the two *H. pylori* flagellar filaments, FlaA and FlaB, within the WT flagellum reveal that they possess distinct glycosylation patterns and form different supercoiled waveforms. These structural differences translate into functional specialization. While both flagellins are required for optimal motility, they antagonistically regulate biofilm initiation: FlaA promotes it, while FlaB suppresses it. Furthermore, FlaB’s suppressive function requires a downstream signaling via the motor cage protein, PilO. Collectively, these results reveal that *H. pylori* evolved a dual-flagellin system with specialized roles to dynamically coordinate motility and surface attachment, facilitating adaptation to the dynamic gastric environment.

## Methods and Materials

### Bacterial strains and growth conditions

*Helicobacter pylori* strains (Table S1) utilized in this work included the wild-type G27 and its derivative mutants, detailed in Table S1. For routine culture, *H. pylori* was grown on either solid Columbia Horse Blood Agar (CHBA; Difco) plates or in liquid Brucella Broth (BB; Difco) supplemented with 10% heat-inactivated fetal bovine serum (HI-FBS; Life Technologies), designated BB10. CHBA media formulations were supplemented with the following additives: 0.2% (w/v) beta-cyclodextrin, 10 µg/ml vancomycin, 5 µg/ml cefsulodin, 2.5 U/ml polymyxin B, 5 µg/ml trimethoprim, and 8 µg/ml amphotericin B. *H. pylori* cultures were incubated at 37°C under microaerobic conditions (5% O₂, 10% CO₂, 85% N₂). Liquid cultures were typically agitated at 200 rpm. Where appropriate, *H. pylori* mutant-selective antibiotics were added to the media at 15 µg/ml kanamycin or 13 µg/ml chloramphenicol. The *Escherichia coli* DH10B strain employed was cultured in LB containing 15 µg/ml kanamycin.

### Mutant construction

Mutants were generated in *H. pylori* G27 using allelic exchange via natural transformation, primarily employing two-step PCR fusion to create replacement cassettes. Unless otherwise specified, mutant alleles were transformed into *H. pylori* G27, selected on CHBA plates with the appropriate antibiotic (chloramphenicol for *cat*, kanamycin for *aphA3*), and verified by sequencing and/or PCR.

The *flaA* mutant (Δ*flaA*::cat) was constructed by replacing the native *flaA* sequence with the *cat* chloramphenicol resistance gene, following a previously described method (27). Construction of the *flaA* allele exemplifies the general two-step PCR fusion protocol: First, upstream (primers XL118/XL119) and downstream (primers XL122/XL123) (Table S2) homology arms flanking the target gene were amplified, alongside the cat gene (primers XL120/XL121; template pCat-mut (74). These three amplicons were fused using primers XL118/XL123. The *flaB* mutant (Δ*flaB::cat*) was created similarly using primers XL124/XL125 (upstream arm), XL128/XL129 (downstream arm), and XL126/XL127 (*cat* gene).

To create double mutants, resistance cassettes were swapped. A Δ*flaA::aphA3* mutant was created by replacing the *cat* cassette in the Δ*flaA::cat* strain with *aphA3*. Homology arms corresponding to the first and last 250 bp of the *cat* gene (primers XL164/XL165 and XL168/XL169; template cat cassette) and the full-length *aphA3* gene (primers XL166/XL167; template *aphA3* cassette) were amplified. These amplicons were fused (primers XL168/XL169), purified (GFX™ PCR DNA and Gel Band Purification Kit), and transformed into the Δ*flaA::cat* mutant, selecting for kanamycin resistance. The Δ*flaB::aphA3* mutant was generated similarly, and it was used to generate Δ*flaA::cat* Δ*flaB::aphA3* by replacing the native *flaA* sequence with the *cat* (Table S1). The Δ*flaA*::*cat* mutant and Δ*flaB::cat* mutant were used as parent strains for replacement of *pilO* with *alpA3,* to construct Δ*flaA*::*ca*t Δ*pilO::aphA3* and Δ*flaB*::*cat* Δ*pilO::aphA3*.

### Semisolid agar migration assay

Motility was assessed using a semisolid agar migration assay. *H. pylori* wild-type (WT) or derivative strains were grown overnight in liquid BB10 medium. Cultures were adjusted to an OD₆₀₀ of 0.15 using fresh BB10 and shaken for 2 hours to ensure full motility. A 2 µl aliquot of each motile culture was spotted onto the center of a semisolid agar plate. The plates consisted of Brucella Broth (BB) supplemented with 2.5% heat-inactivated fetal bovine serum (HI-FBS) and 0.35% (w/v) agar (Bacto). Plates were incubated under standard microaerobic conditions (37°C, 5% O_2_, 10% CO_2_, 85% N_2_) for 3 days. Images for cultures incubated 3 days were acquired using a Gel Doc XR+ Gel Documentation System (Bio-Rad). The diameter of the migration halo was measured using the measurement tool in ImageJ software (https://imagej.nih.gov/ij/index.html).

### Swimming assay

*H. pylori* strains were cultured overnight in BB10 liquid medium under standard microaerobic conditions with shaking (200 rpm, 37°C). Cultures were adjusted to an OD₆₀₀ of 0.15 using fresh BB10 and incubated for an additional 2 hours to ensure motility. For swimming analysis, 70 μl aliquots were transferred to concave-well slides, covered with coverslips, and immediately observed under phase-contrast microscopy (Nikon ECLIPSE E600, 40× objective). Videos (20 seconds duration) were recorded using a Hamamatsu C7472-95 digital camera. Cell trajectories were analyzed using the TrackMate plugin in ImageJ to calculate swimming speeds. Reversal frequencies were manually determined from trajectory plots.

### Crystal violet assay

*H. pylori* G27 wild-type and mutant strains were cultured overnight in BB10 liquid medium. Cultures were adjusted to OD_600_ = 0.15 in BB10, and 200 μl aliquots were transferred to wells of a sterile 96-well polystyrene microtiter plate (Costar #3596). Plates were incubated statically under microaerobic conditions at 37°C for 1-3 days. After incubation, media was removed by pipetting and wells were washed twice with 200 μl phosphate-buffered saline (PBS). Biofilms were stained with 200 μl of 0.1% (w/v) crystal violet for 10 minutes at room temperature. Following dye removal, wells were washed twice with PBS. Bound dye was solubilized with 200 μl of 90% (v/v) ethanol, and biofilm biomass was quantified by measuring absorbance at 595 nm using a microplate reader.

### Scanning Electron Microscopy and Flagella Analysis

*H. pylori* G27 WT or mutant cells from BB10 liquid cultures were adsorbed onto 12-mm round glass coverslips (Costar) for 5-10 minutes. Attached cells were fixed with 2.5% (w/v) glutaraldehyde for 1 hour at room temperature. Samples underwent ethanol dehydration through a graded series (25%, 50%, 75%, 90%, and two changes of 100% ethanol; 10 minutes per step). Dehydrated specimens were critical point dried (Balzers Union 342/11 120B), sputter-coated with ≈20 nm gold (Hammer IV, Technic Inc.), and imaged using an FEI Quanta 3D dual-beam SEM at 5 kV and 6.7 pA. Biofilm-grown cells on coverslips underwent identical fixation, dehydration, drying, coating, and imaging procedures. Flagella number and length were quantified from the resulting micrographs.

### Confocal assay

Biofilm samples were prepared following the same initial culture protocol as the biofilm formation assay (overnight growth in BB10, adjustment to OD_600_ = 0.15). 300 µL aliquots were transferred to wells of µ-Slide 8-well glass-bottom chamber slides (ibidi, Germany), each containing a vertically positioned coverslip. Slides were incubated statically under microaerobic conditions at 37°C for 1 or 3 days. After incubation, planktonic cells and media were removed. Biofilms were washed three times with PBS to eliminate unattached material. 400 µL PBS was added to each well, and biofilm architecture was imaged using a Zeiss 880 confocal microscope with 488-nm laser excitation.

### Cryo-EM of *H. pylori* flagella

Cells were cultured in two 250 mL flasks, each containing 25 mL of medium. The 50 mL total culture was centrifuged at 8,000 rpm for 10 min. The cell pellet was resuspended in phosphate-buffered saline (PBS; pH 7.2) at a ratio of 1 mL PBS per 4 mL of original culture volume. Flagellar filaments were sheared from the cells by vortexing the suspension at maximum speed for 15 min. Cell debris and intact cells were removed by centrifugation at 10,000 × g for 10 min. The supernatant containing sheared flagella was used for cryo-EM imaging.

The flagella sample (4.5 μL) was applied to glow-discharged lacey carbon grids and then plunge-frozen using an EM GP2 Plunge Freezer (Leica). The cryo-EM micrographs were collected on a 300 keV Titan Krios with a K3 camera at ∼1.1 Å per pixel and a total dose of 50 e-/Å2. The cryo-EM workflow was initiated with patch motion corrections and CTF estimations in cryoSPARC (75–77). Following this, particles were auto-picked using the ‘Filament Tracer’ function with a shift of 14 pixel between adjacent boxes. For reconstruction of the FlaB from WT flagella, filaments without a clearly identifiable hook were manually removed. For asymmetric reconstruction, adjacent particles within 50 Å were removed. All particles were subsequently 2D classified with multiple rounds, and all particles in bad 2D averages were removed. The canonical bacterial flagellum symmetry was confirmed by looking at the average power spectrum of aligned raw particles (Fig. S4). For refinement with helical symmetry, 3D reconstruction was performed using ‘Helical Refine’ first, ‘Local CTF refinement’ next, and another round of ‘Helical Refine’ using the CTF-refined particles. For asymmetrical reconstruction, 640-pixel boxes were re-exacted after ‘Helical Refine’, followed by ‘Homogenous Refinement’ to relax the helical symmetry. After that, we performed another round of ‘Local CTF refinement’ and ‘Homogenous Refinement’. The resolution of each reconstruction was estimated by Map:Map FSC, Model:Map FSC, and d99 (78). Maps used in the resolution estimation were sharpened using Local Filter available in cryoSPARC. The statistics are listed in the Table S3.

### Model building of flagellar filaments

Regarding the starting reference, the AlphaFold3 predicted structure of a single flagellin subunit was initially docked into the cryo-EM map. Domains were docked individually since AlphaFold predictions for domain-domain orientation are frequently imprecise. The subunit and glycosylation residues were then manually adjusted and refined in Coot (79). Specifically, previously reported pseudaminic acids (46) were added at sites where putative monosaccharide densities were observed extending from serine or threonine residues. Bond and angle restraints for Pse were added based on high-resolution X-ray structure (80) (PDB ID: 5JSD). For helical averaged maps, a filamentous model can be extended from the single flagellin.

For curved flagellar map, the refined single flagellin was docked into the other ten protofilaments within the supercoiled flagellar filament map and subjected to real-space refinement via PHENIX (81). The filament model’s quality was assessed using MolProbity (82), and the refinement statistics for all three flagellar filaments are detailed in Table S3. Cryo-EM map and model visualization were primarily done in ChimeraX (83).

For the comparative analysis of FlaA glycosylation between wild-type and Δ*flaB* mutant, the respective cryo-EM maps were sharpened using the same B-factor, and their thresholds were normalized based on the average density of the conserved D0/D1 core domains to enable a direct comparison of glycan densities.

### Quantitative RT-PCR (qRT-PCR)

qRT-PCR was performed on *flaA* and *flaB* genes. Total RNA was extracted from three independent biological replicates. For each reaction, 1 µg of total RNA was reverse-transcribed into cDNA using the LunaScript^TM^ RT SuperMix Kit (New England Biolabs). qPCR reactions were prepared with Luna Universal qPCR Master Mix (New England Biolabs) and run on a Connect Thermal Cycler (Bio-Rad) under the following cycling conditions: initial denaturation at 95°C for 2 minutes; 39 cycles of 95°C for 30 seconds, 60°C for 30 seconds, and 72°C for 30 seconds. Gene-specific primers were designed using NCBI Primer-BLAST. The 16S rRNA gene was used as an endogenous control for normalization, as its expression was stable across conditions. Relative gene expression was calculated using the 2 ^−ΔΔCT^ method. All qPCR analyses were performed with three biological replicates and two technical replicates per sample.

## Acknowledgements

We thank all members of the Ottemann Lab for helpful discussions and comments on the paper. We thank Benjamin Abrams at the UCSC Life Sciences Microscopy Center for technical support with confocal microscopy (RRID: SCR_021135) for help with biofilm imaging. This research was, in part, supported by the National Cancer Institute’s National Cryo-EM Facility at the Frederick National Laboratory for Cancer Research under contract 75N91019D00024. Electron microscopy screening was carried out in the UAB Cryo-EM Facility, supported by the Institutional Research Core Program and O’Neal Comprehensive Cancer Center (NIH grant P30 CA013148), with additional funding from NIH grant S10 OD024978. We are grateful to Dr. James Kizziah, and Dr. Tara Fox for assisting with the screening or data collection. The described project was supported by the National Institute of Allergy and Infectious Disease (NIAID) grants RO1 AI116946 and AI164682 to K.M.O., Guangxi Science and Technology Program 2025JJA130146, National Natural Science Foundation of China (NSFC) grant 32500161, and Ba-Gui Young Scientist Support Program of Guangxi to X.L. The cryo-EM work in F.W. laboratory was supported by NIH grant R35GM159704.

## Supplementary Material

**Table S1.**
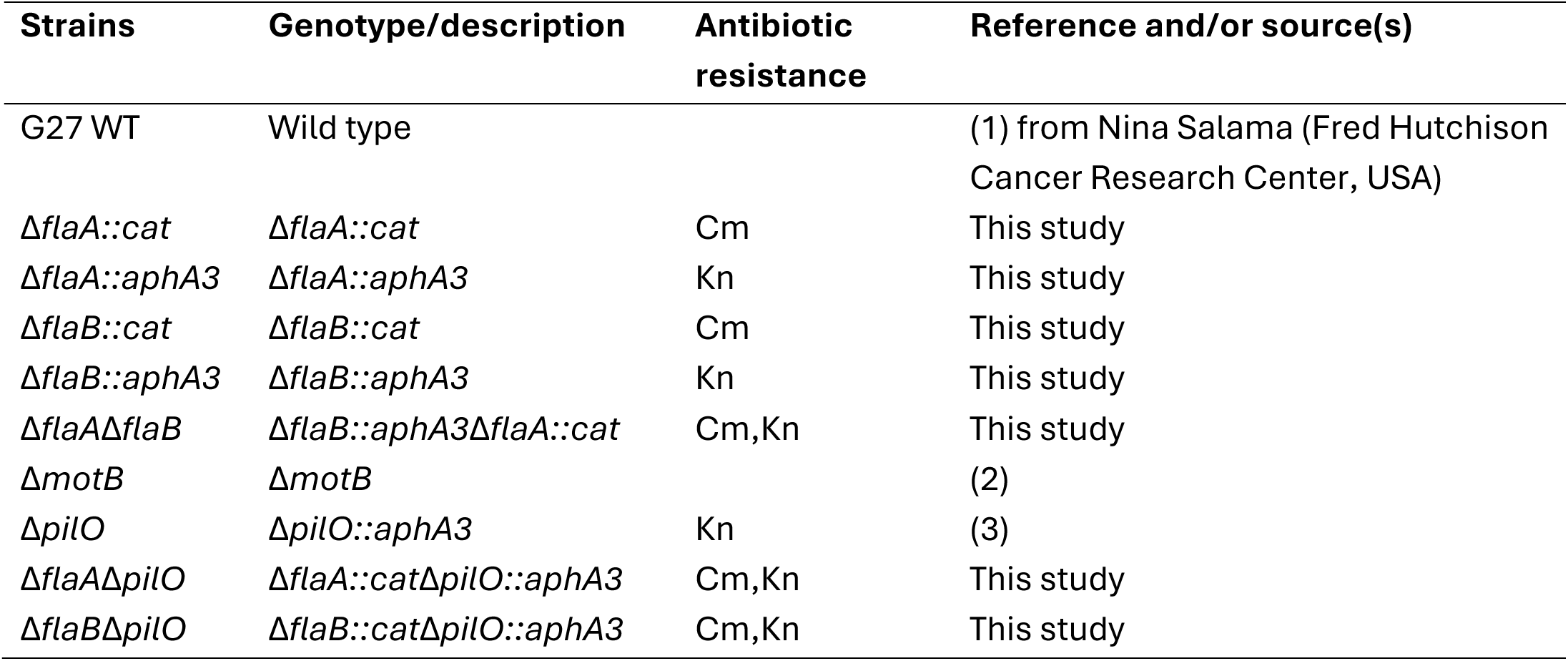
Strains used in this study.

| Strains | Genotype/description | Antibiotic resistance | Reference and/or source(s) |
| --- | --- | --- | --- |
| G27 WT | Wild type |  | (1) from Nina Salama (Fred Hutchison Cancer Research Center, USA) |
| *ΔflaA::cat* | *ΔflaA::cat* | Cm | This study |
| *ΔflaA::aphA3* | *ΔflaA::aphA3* | Kn | This study |
| *ΔflaB::cat* | *ΔflaB::cat* | Cm | This study |
| *ΔflaB::aphA3* | *ΔflaB::aphA3* | Kn | This study |
| *ΔflaAΔflaB* | *ΔflaB::aphA3ΔflaA::cat* | Cm,Kn | This study |
| *ΔmotB* | *ΔmotB* |  | (2) |
| *ΔpilO* | *ΔpilO::aphA3* | Kn | (3) |
| *ΔflaAΔpilO* | *ΔflaA::catΔpilO::aphA3* | Cm,Kn | This study |
| *ΔflaBΔpilO* | *ΔflaB::catΔpilO::aphA3* | Cm,Kn | This study |

**Table S2.**
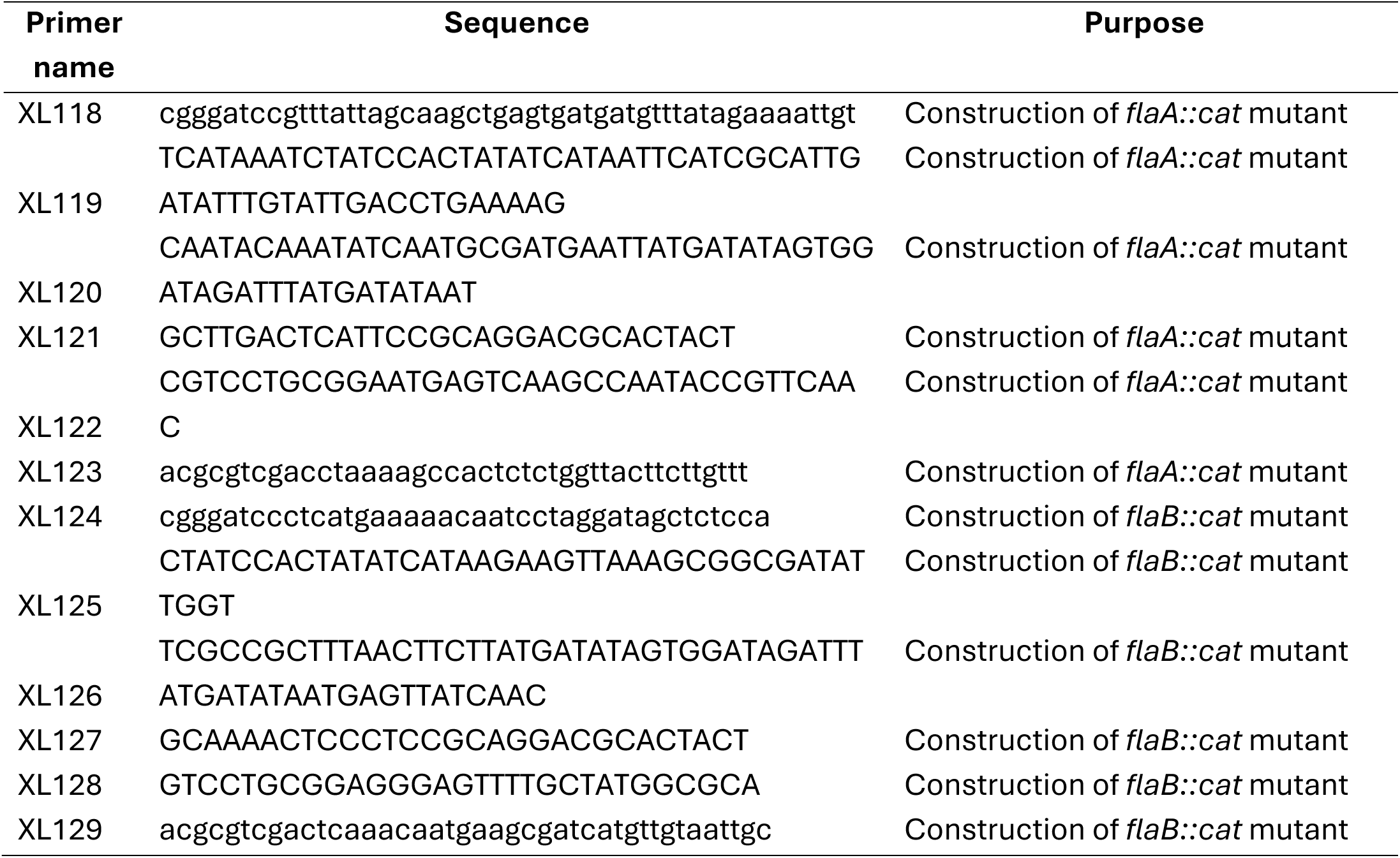
Primers used in this study.

**Table S3.** Cryo-EM and Refinement Statistics of flagellar filaments.

| Parameter | wildtype (FlaA region) | wildtype (FlaB region) | FlaA ($\Delta flaB$) | FlaB ($\Delta flaA$) |
| --- | --- | --- | --- | --- |
| **Data collection and processing** |  |  |  |  |
| Voltage (kV) | 300 | 300 | 300 | 300 |
| Electron exposure ($e^- \text{Å}^{-2}$) | 50 | 50 | 50 | 50 |
| Pixel size (Å) | 1.09 | 1.09 | 1.07 | 1.07 |
| Particle images (n) | 2,351,759 | 222,264 | 235,055 | 66,393 |
| Shift (pixel) | 6 | 6 | 55 | 55 |
| **Helical symmetry (if applied)** |  |  |  |  |
| Point group | C1 | C1 | C1 | C1 |
| Helical rise (Å) | 4.77 | 4.78 | N.A. | N.A. |
| Helical twist (°) | 65.40 | 65.40 | N.A. | N.A. |
| **Map resolution (Å)** |  |  |  |  |
| Map:map FSC (0.143) | 2.8 | 3.2 | 3.0 | 3.4 |
| Model:map FSC (0.5) | 2.8 | 3.2 | 3.1 | 3.8 |
| d<sub>99</sub> | 3.1 | 3.8 | 3.3 | 3.8 |
| **Refinement and Model validation** |  |  |  |  |
| Ramachandran Favored | 98.5 | 93.32 | 98.52 | 94.68 |
| Ramachandran Outliers | 0.00 | 0.00 | 0.00 | 0.04 |
| Real Space CC | 0.90 | 0.86 | 0.80 | 0.81 |
| Clashscore | 15.21 | 7.70 | 7.3 | 10.9 |
| Bonds RMSD, length (Å) | 0.002 | 0.003 | 0.002 | 0.003 |
| Bonds RMSD, angles (°) | 0.450 | 0.505 | 0.453 | 0.506 |
| **Deposition ID** |  |  |  |  |
| PDB (model) | 9XZ0 | 9XZ3 | 9XZ4 | 9XZ7 |
| EMDB (map) | EMD-72345 | EMD-72347 | EMD-72348 | EMD-72351 |

**Figure S1.**
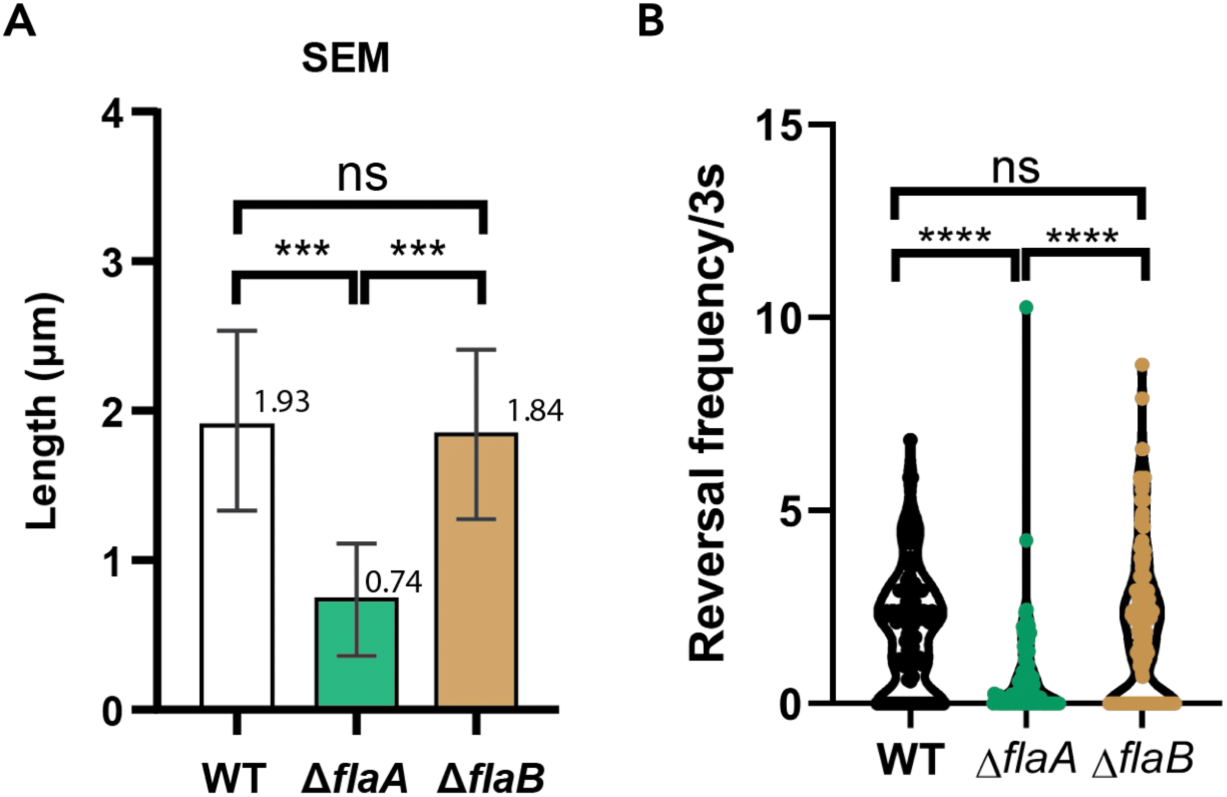
Quantification of flagellar length, and frequency of changing swimming directions in *H. pylori* WT, Δ*flaA*, and Δ*flaB* strains. (A) Flagellar lengths were measured from scanning electron micrographs of wild-type (WT), *flaA* deletion mutant (Δ*flaA*), and *flaB* deletion mutant (Δ*flaB*) cells. Bars represent the mean ± standard error of the mean. Statistical significance was determined by Student’s t test; ***p < 0.001, ns = not significant. The discrepancy between lengths measured by SEM and TEM likely reflects the higher accuracy of TEM for the purpose of measuring cell filament lengths. With negative-stain TEM, it is easier to identify the start of a filament, in ideal cases the course of a filament can be visualized as it passes through the projection of the cell body. (B) The frequency of changing swimming directions of *H. pylori* WT, Δ*flaA*, and Δ*flaB* cells. The data are presented as violin plots computed from the data using a standard univariate gaussian kernel density estimate. The asterisks indicate a significant difference between strains according to one-way ANOVA with Tukey’s correction (**, p<0.0015; ****, p<0.0001).

**Figure S2.**
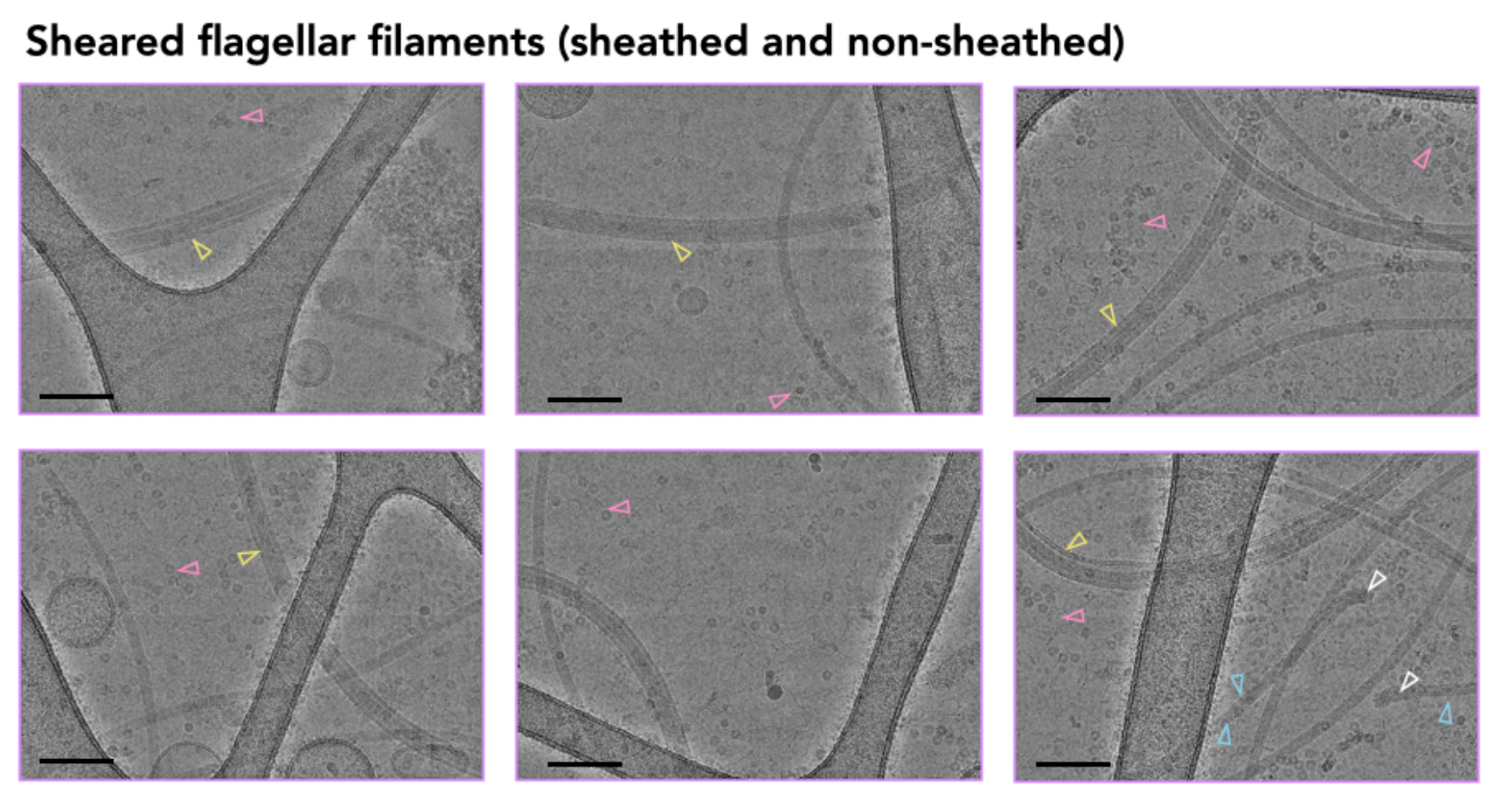
Cryo-EM micrographs of sheared wildtype *H. pylori* flagellar filaments. The observed flagellar hook is indicated by a white arrowhead. FlaB filaments are indicated by blue arrowheads. The flagellar filaments with visible sheath are indicated by yellow arrowheads. Observed ureases are indicated by a pink arrowhead. Scale bar: 100 nm.

**Figure S3.**
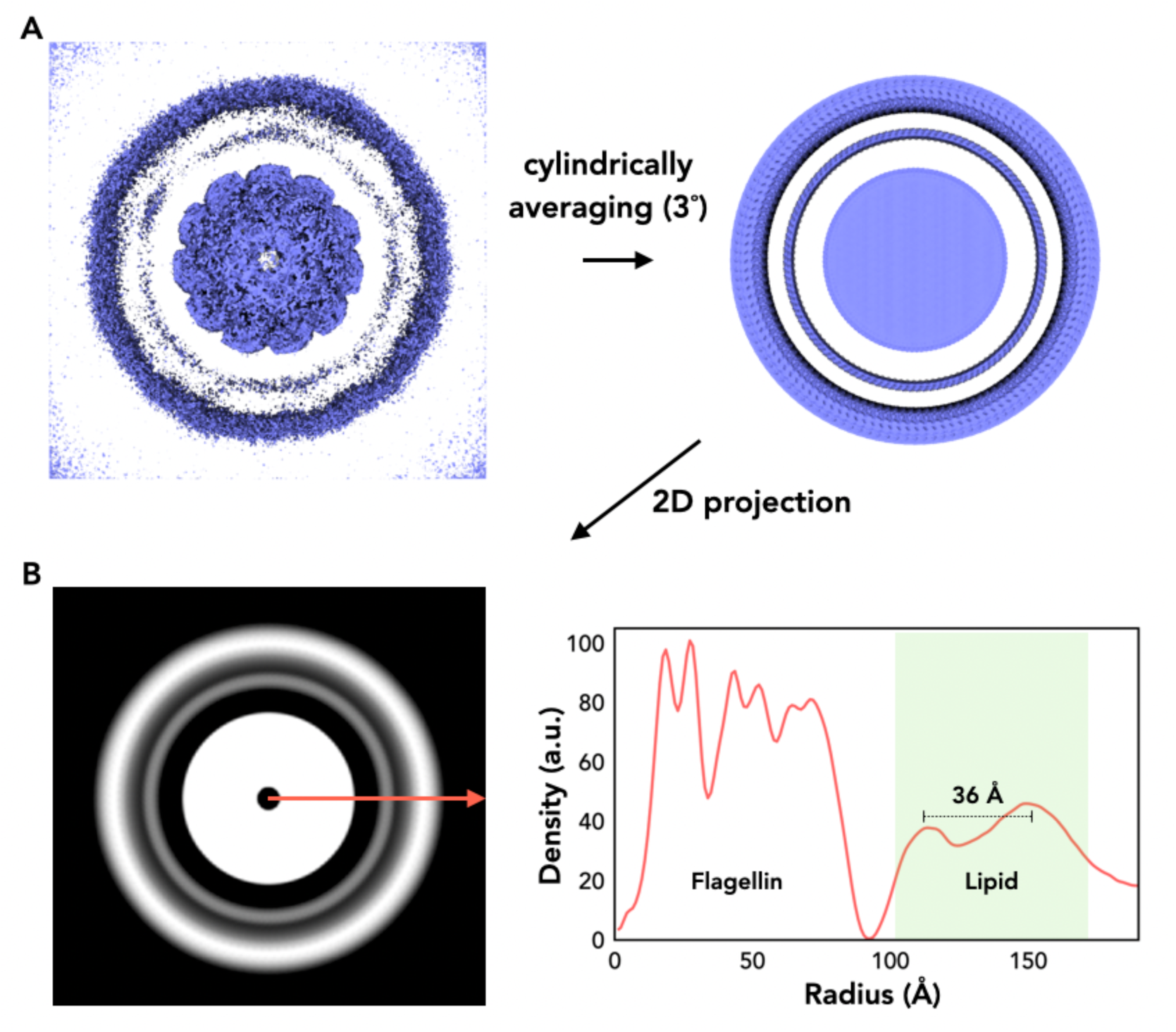
The cryo-EM density of the H. pylori flagellar sheath exhibits lipid bilayer properties. (A) The helically averaged reconstruction of the sheath flagella (left). On the right is the cylindrically averaged volume, generated by rotating the volume on the left in 3-degree increments and summing the resulting volumes. (B) The projection of the cylindrically averaged volume (left) and its radial density profile (right).

**Figure S4.**
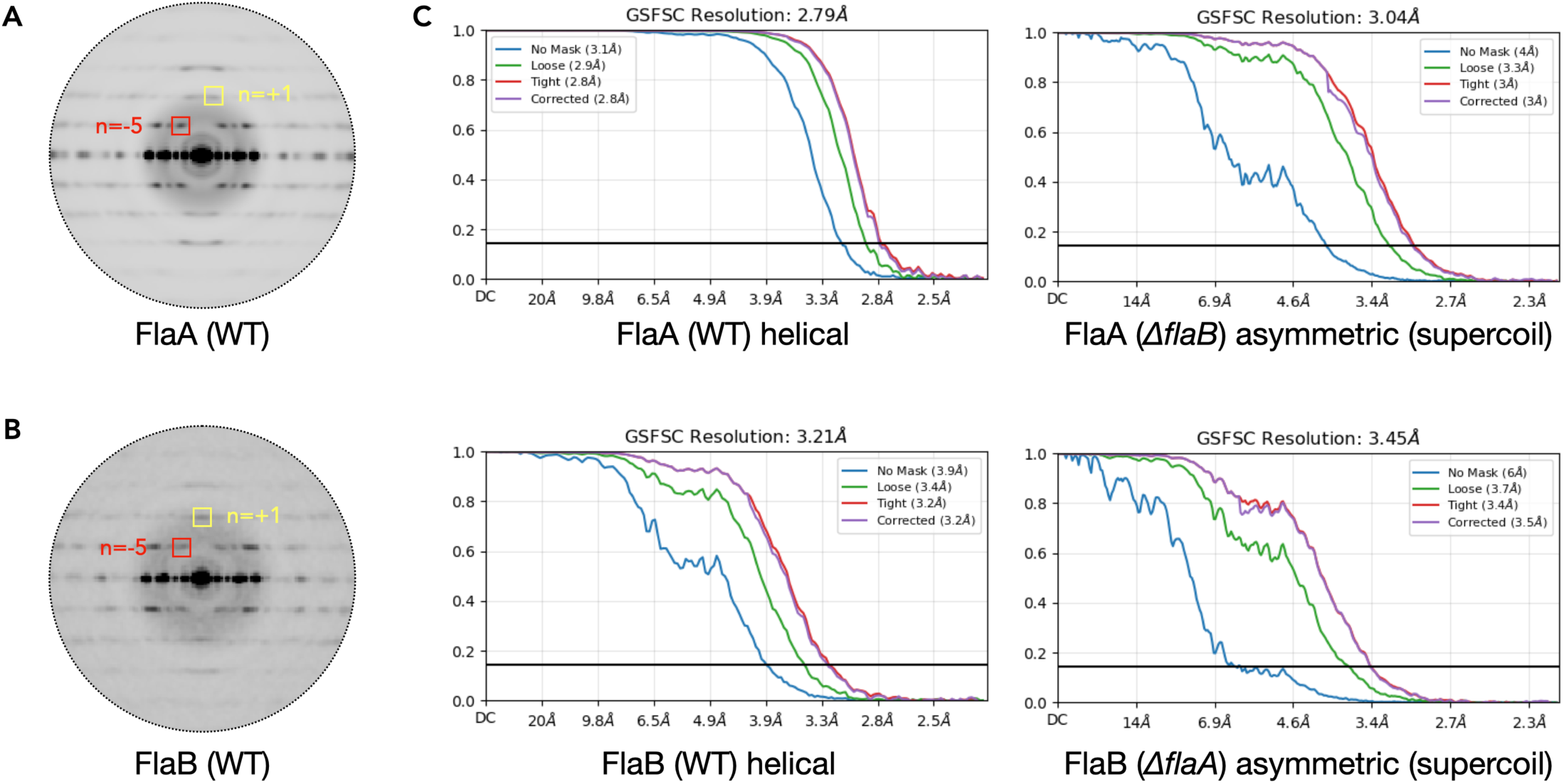
Fourier Shell Correlation (FSC) calculations and average power spectra of flagellar filaments. (A-B) Average power spectra of particles behind 2D class average of FlaA region in wildtype flagellar filaments (A), and FlaB region in wildtype flagellar filaments (B). The Bessel orders of the layer lines from canonical flagellar are labeled. (C) The map:map “gold-standard FSC” curves for 360-pixel box helical reconstruction and 640-pixel box asymmetrical reconstruction of different flagellar filaments.

**Figure S5.**
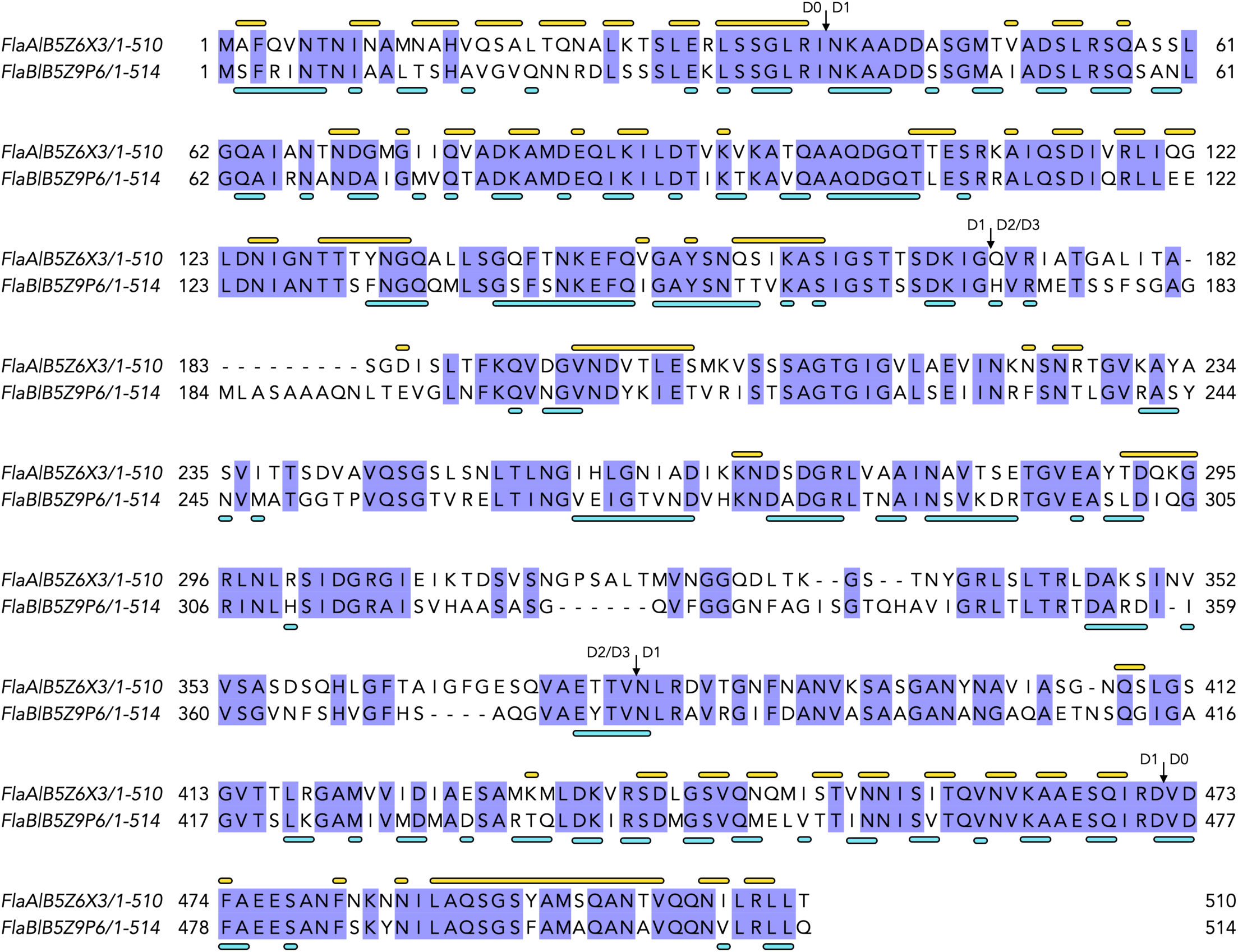
Sequence alignment of FlaA and FlaB in *H. pylori*. In this sequence alignment, identical residues between FlaA and FlaB are highlighted in blue. The residues involved in the left-handed 5-start and 11-start interfaces are indicated by yellow and cyan bars placed above and below the sequences, respectively. The boundaries separating the D0, D1, and D2/D3 domains are also marked.

**Figure S6.**
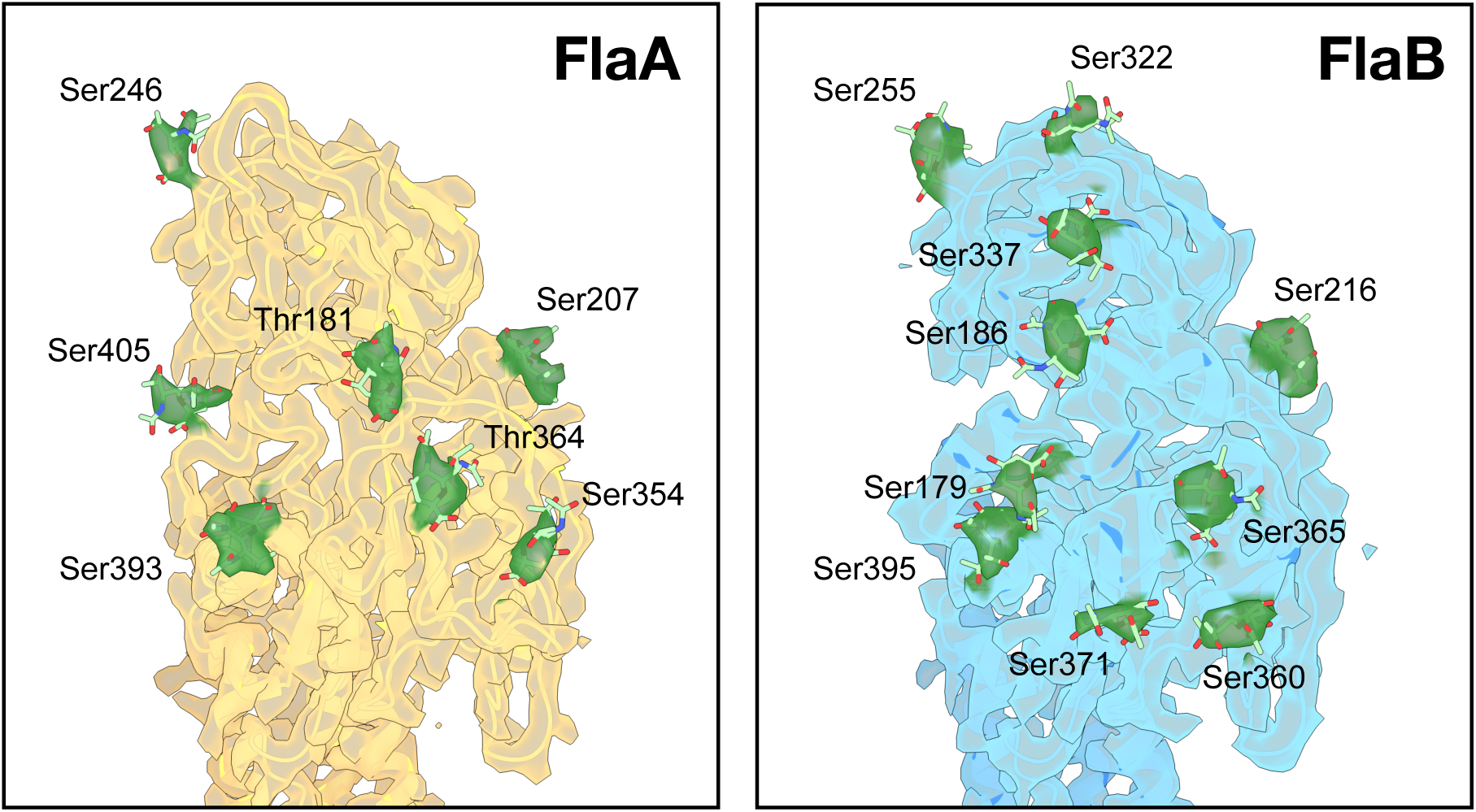
Cryo-EM maps & models of FlaA and FlaB with putative glycans. Cryo-EM maps reconstructed with helical symmetry are shown for FlaA and FlaB from the wild-type flagella filament. Masks for a single-subunit were applied and map voxels within 2.0 Å of a glycan in the refined glycosylated flagellin models are colored green. In total, FlaA contains 7 glycosylated residues (5 serine and 2 threonine), while FlaB contains 10 glycosylated serine residues.

**Figure S7.**
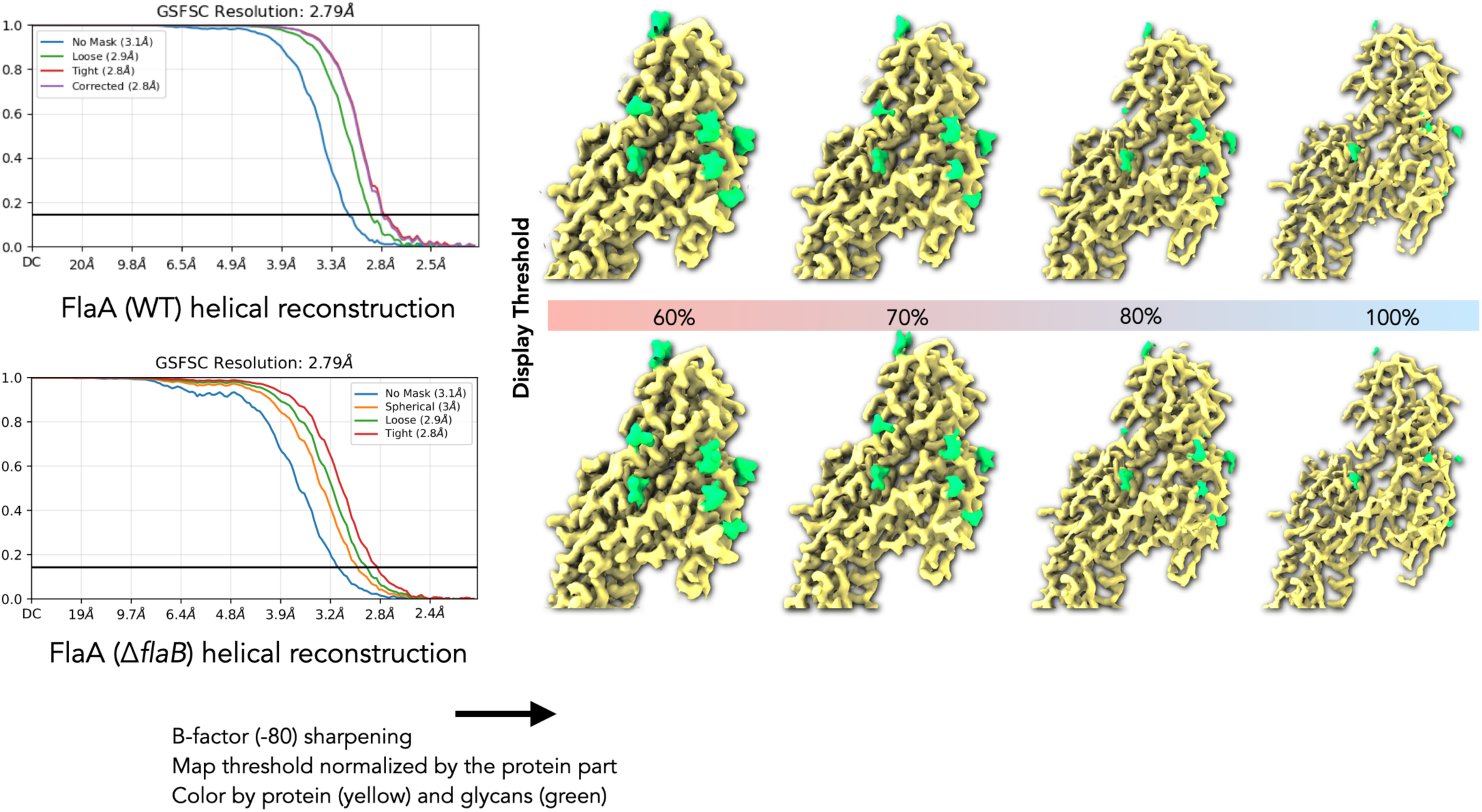
Comparative analysis of FlaA glycosylation in wild-type and FlaA-only (Δ*flaB*) filaments. Left: The map:map “gold-standard FSC” curves for 360-pixel box helical reconstructions of wild-type FlaA and FlaA-only (Δ*flaB* mutant) filaments. Right: Both maps were sharpened with an identical B-factor, and normalized based on the protein density of the D0/D1 domains. Zoom-in comparison of the FlaA protein (yellow surface) with associated glycan densities (green surface) shown at the different display threshold.

**Figure S8.**
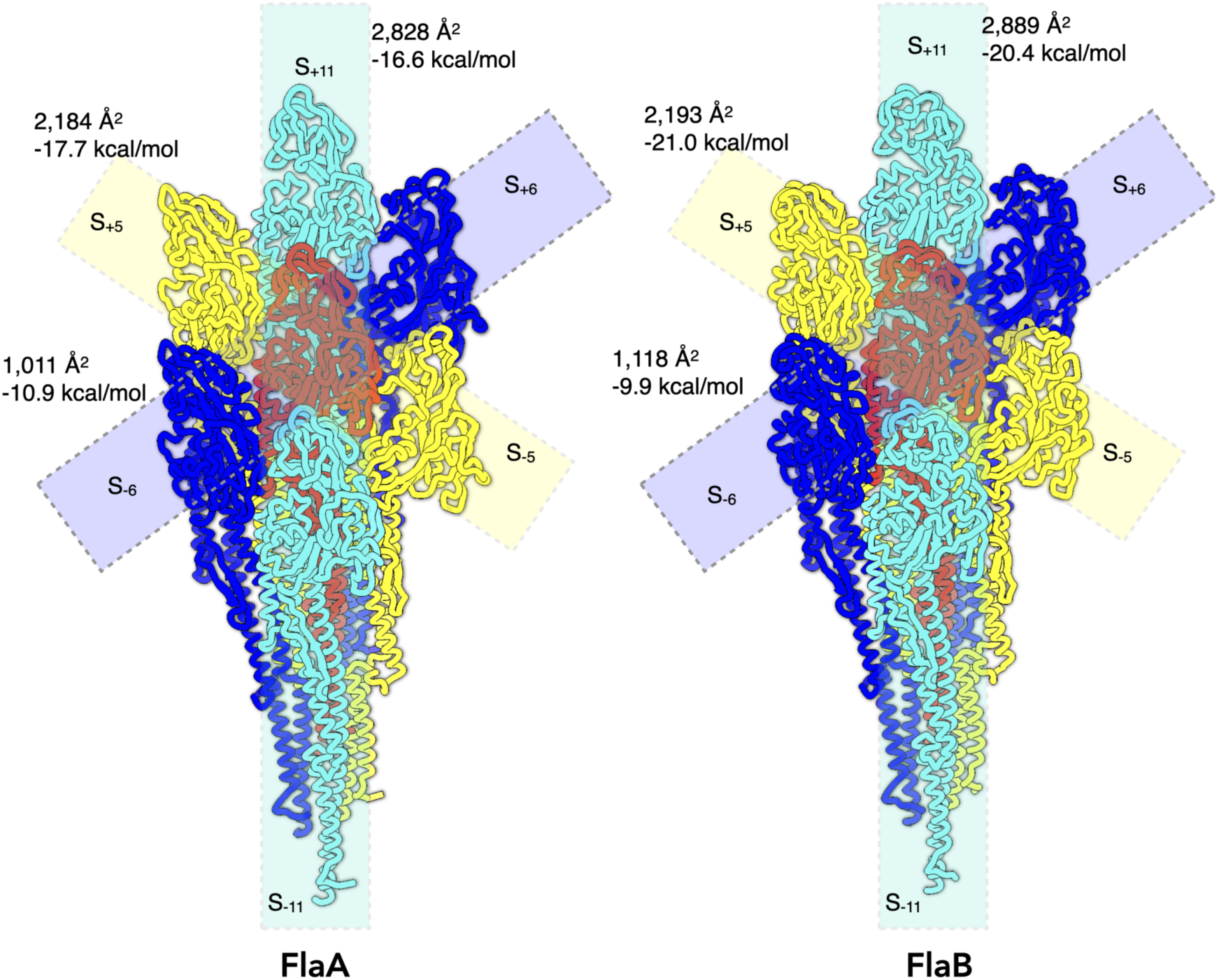
PISA Interface Analysis of FlaA and FlaB Filaments. PISA analysis was performed using helical models of FlaA and FlaB filaments. The major interfaces between flagellins (11-start, 5-start, and 6-start) are indicated. The estimated interface area, and solvation free energy gain upon interface formation are also shown for each interface between two flagellins.

**Figure S9.**
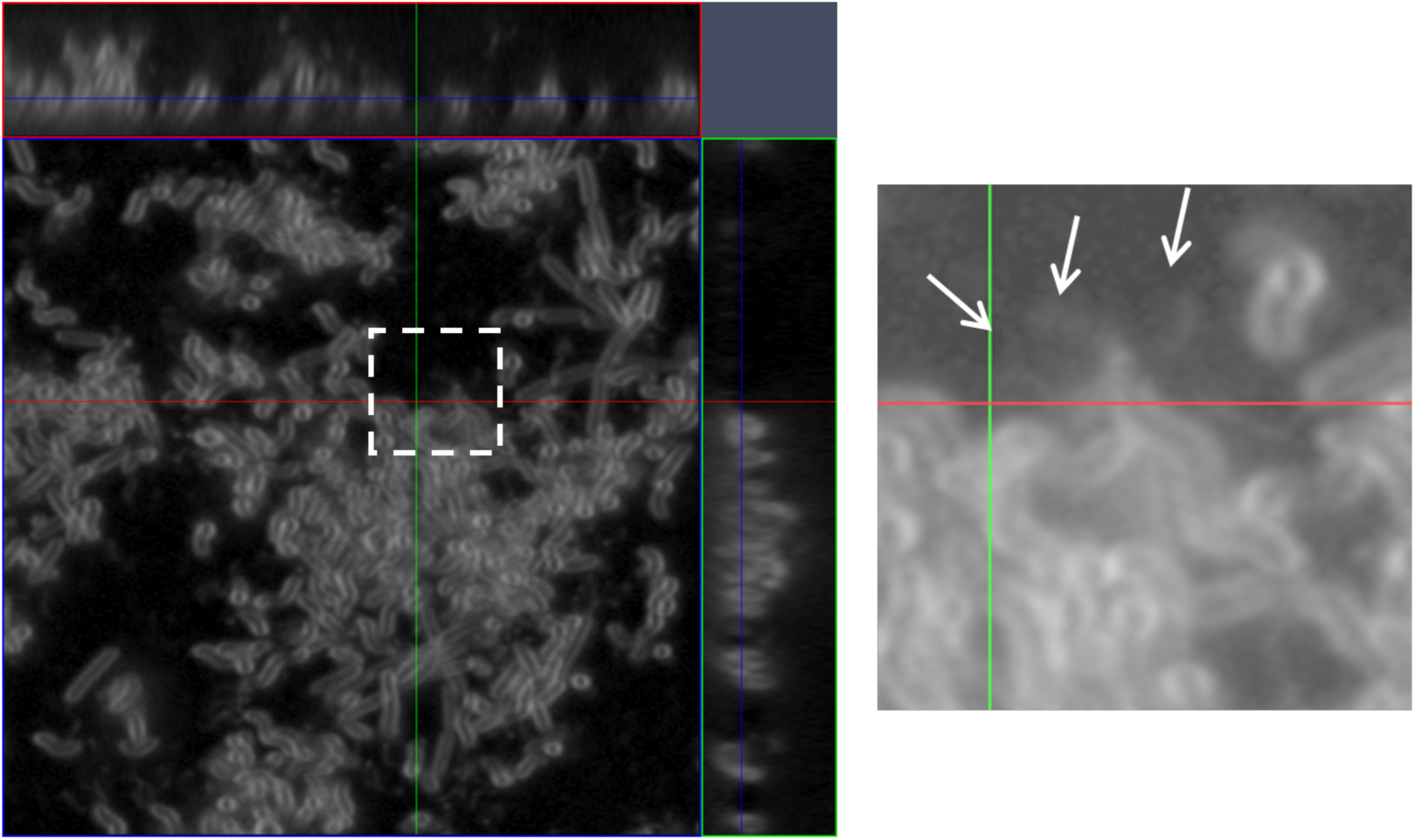
Visualization of flagellar filaments in *H. pylori* strains by confocal microscopy. Flagellar presence and membrane connections (arrowheads) in wild-type (WT) cells were confirmed using the lipophilic dye FM4-64.

**Figure S10.**
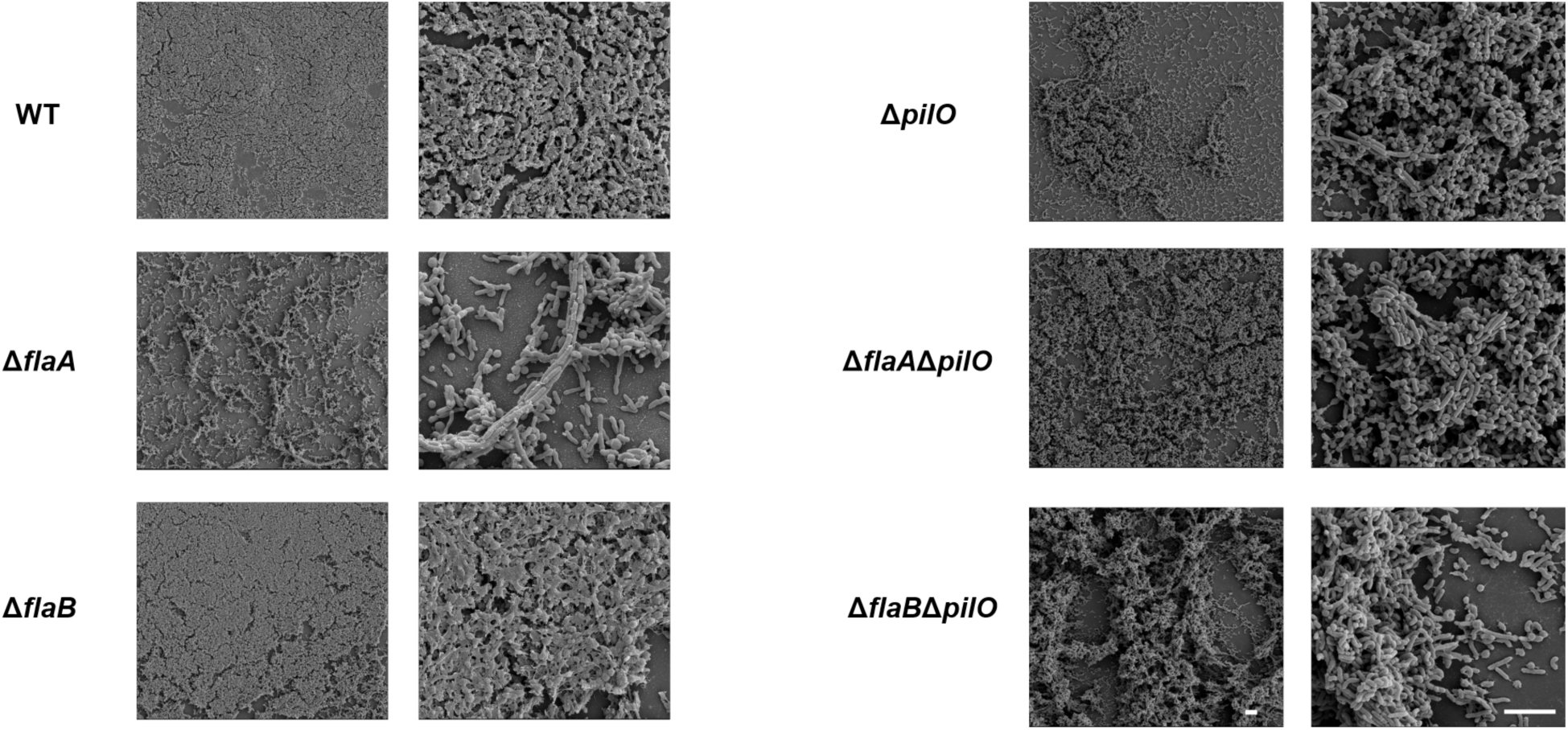
Biofilm formation of *H. pylori* G27 WT and derivative strains. Scanning electron micrographs (SEM) of *H. pylori* G27 Δ*flaA*Δ*flaB* and Δ*motB* cells after culturing on glass surface for 3 days (B). WT and Δ*flaB* mutants form thick biofilms with extensive cell clustering and extracellular material. Δ*flaA* mutants develop rough biofilms characterized by rope-like structures formed through direct cell-cell contact (lacking flagellar bridging). Scale bars: 5 µm.

**Figure S11.**
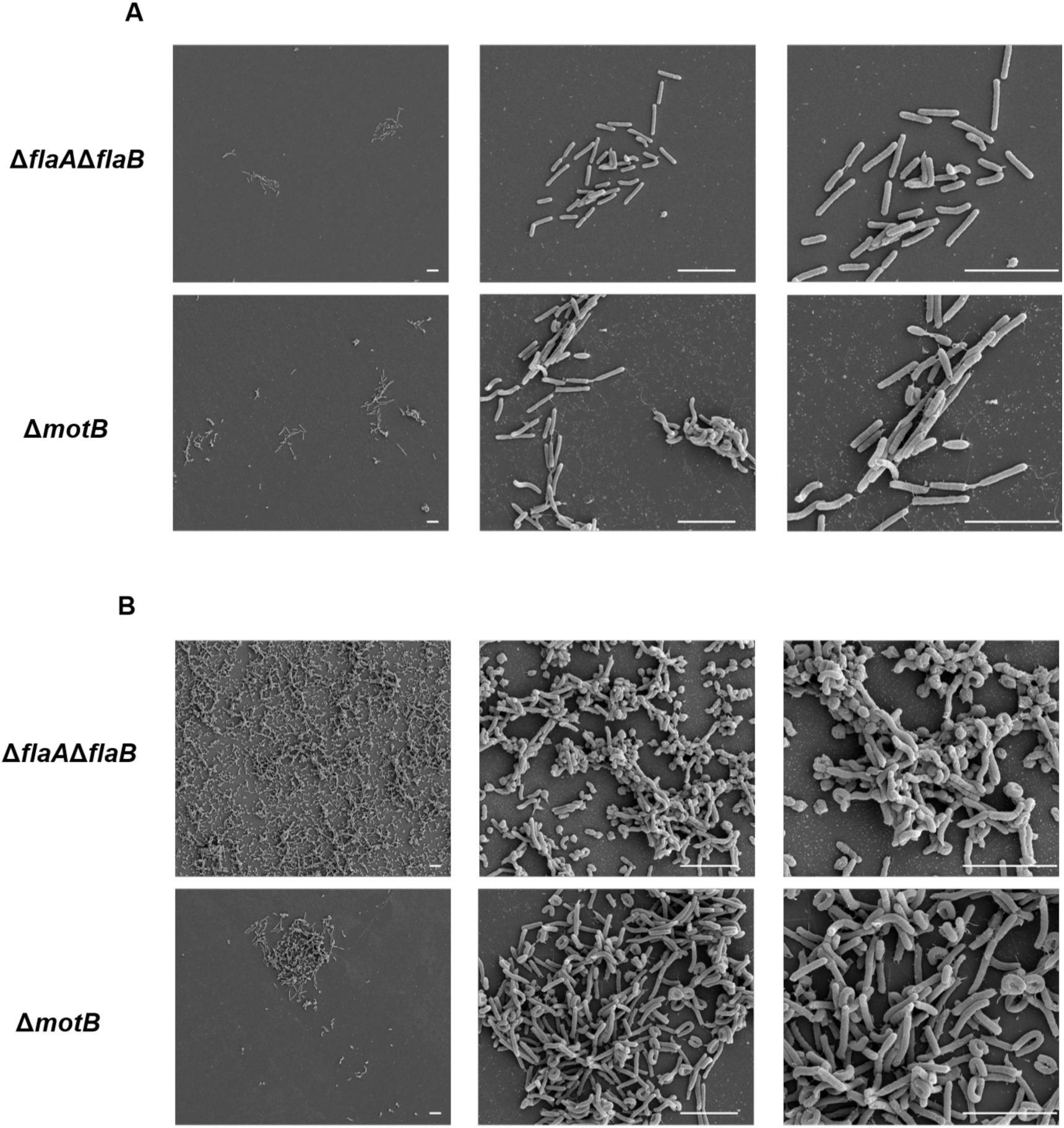
FlaAB and MotB are required for biofilm cluster formation. Scanning electron micrographs (SEM) of *H. pylori* G27 Δ*flaA*Δ*flaB* and Δ*motB* cells after culturing on glass surface for 1 day (A) and 3 days (B). (A) After 1 day, the afflagelate Δ*flaA*Δ*flaB* mutant and non-motile Δ*motB* mutant have low biofilm capacity. (A) After 3 days, only Δ*flaA*Δ*flaB* and Δ*motB* exhibited loose biofilm formation. The scale bar indicates 5 μm.

**Figure S12.**
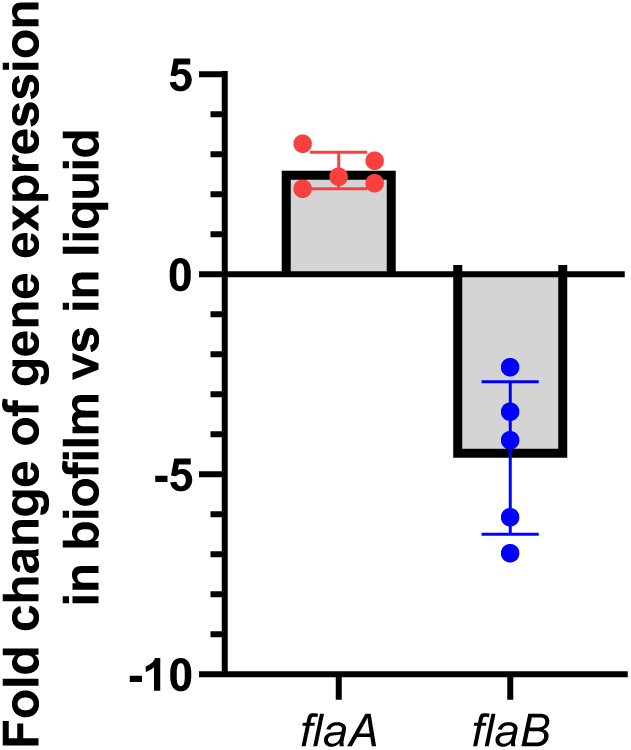
The changes of *flaA* and *flaB* in biofilms versus in liquid. mRNA was isolated from *H. pylori* G27 WT cells grown in biofilm or planktonic conditions, reverse transcribed to cDNA, and analyzed by RT-qPCR. Expression of *flaA* was increased in biofilms, while expression of *flaB* was decreased, relative to planktonic cultures.

